# NF-κB modifies the mammalian circadian clock through interaction with the core clock protein BMAL1

**DOI:** 10.1101/2020.09.06.285254

**Authors:** Yang Shen, Wei Wang, Mehari Endale, Lauren J. Francey, Rachel L. Harold, David W. Hammers, Zhiguang Huo, Carrie L. Partch, John B. Hogenesch, Zhao-Hui Wu, Andrew C. Liu

**Affiliations:** Department of Physiology and Functional Genomics, University of Florida College of Medicine; Department of Radiation Oncology and Center for Cancer Research, University of Tennessee Health Science Center; Divisions of Human Genetics and Immunobiology, Cincinnati Children’s Hospital Medical Center; Department of Chemistry and Biochemistry, University of California Santa Cruz; Department of Pharmacology and Therapeutics, University of Florida College of Medicine; Department of Biostatistics, College of Public Health & Health Professions, University of Florida College of Medicine

**Keywords:** Circadian clock, inflammation, NF-κB, suprachiasmatic nucleus (SCN), BMAL1

## Abstract

In mammals, the circadian clock coordinates various cell physiological processes, including the inflammatory response. Recent studies suggested a crosstalk between these two pathways. However, the mechanism of how inflammation affects the circadian clock is not well understood. Here, we investigated the role of the proinflammatory transcription factor NF-κB in regulating clock function. Using a combination of genetic and pharmacological approaches, we show that perturbation of the canonical NF-κB subunit *RELA* in the U2OS cellular model altered core clock gene expression. While RELA activation shortened period length and dampened amplitude in these cells, its inhibition lengthened period length and caused amplitude phenotypes. NF-κB perturbation also altered circadian rhythms in the master suprachiasmatic nucleus (SCN) clock and locomotor activity. We show that RELA, like the clock repressor CRY1, potently repressed the transcriptional activity of BMAL1/CLOCK at the circadian E-box cis-element. Biochemical and biophysical analysis showed that RELA competes with coactivator CBP/p300 for binding to the transactivation domain of BMAL1. This mechanism is further supported by chromatin immunoprecipitation analysis showing that the binding sites of RELA, BMAL1 and CLOCK converge on the E-boxes of clock genes. Taken together, these data support a significant role for NF-κB in directly regulating circadian clock function and highlight mutual regulation between the circadian and inflammatory pathways.

## INTRODUCTION

Endogenous circadian clocks allow organisms to coordinate behavior, physiology and metabolism to align with the external light-dark cycle (Bass, 2016). Circadian clocks are present in virtually all cells of the body and are coordinated by the master clock in the suprachiasmatic nucleus (SCN) of the hypothalamus (Liu et al., 2007a; Mohawk et al., 2012). The SCN receives photic inputs and relays the light/dark information to peripheral tissues via neural and endocrine signals. In this manner, the SCN coordinates the peripheral oscillators into a coherent timing system (Mohawk et al., 2012). The circadian clock regulates physiological functions in various tissues in mammals, including the immune and inflammatory responses (Labrecque and Cermakian, 2015; Richards and Gumz, 2013; Skarke et al., 2017). Given the physiological importance of circadian timing, it is not surprising that its disruption is associated with a variety of pathological conditions and disease states, including sleep and neuropsychiatric disorders, metabolic syndrome, cardiovascular diseases, and cancer (Takahashi et al., 2008; Zandra E. Walton et al., 2018; Wulff et al., 2010).

In individual cells, the molecular clock is based on a transcriptional/translational negative feedback mechanism, in which activators drive the expression of their own repressors (Takahashi, 2017). More specifically in mammals, the bHLH-PAS domain-containing transcriptional activators BMAL1 and CLOCK form a heterodimeric complex, which binds to the circadian E-box cis-elements to activate transcription of target genes, including Period (*PER1, 2, 3*) and Cryptochrome (*CRY1, 2*). The PER/CRY proteins form their own complexes, and upon translocation to the nucleus, suppress BMAL1/CLOCK activity. This core loop interacts with at least two other feedback loops mediated by the circadian D-box and RORE cis-elements. These loops serve to stabilize the core loop and increase system robustness. The molecular clock regulates thousands of output genes that govern cell physiology, largely in a tissuespecific manner (Takahashi, 2017; Zhang et al., 2014).

While the core clock mechanism is well understood and significant progress has been made in characterizing clock outputs, there exist additional clock components and modifiers. In an effort to identify additional clock factors, we carried out a genome-wide RNAi screen in a human U2OS cell model and identified hundreds of genes whose knockdown impacted clock function (Zhang et al., 2009). These genes represent many cellular processes and signaling pathways that serve as inputs and link cellular functions to the clock (Morris et al., 2020; Zhang et al., 2009). These input pathways reflect the extensive interplay between the clock and cell physiology. Several emerging examples include nutrient/energy levels and redox stress (Morris et al., 2020; Reinke and Asher, 2019). In this study, we focused on how the pro-inflammatory nuclear factor-kappa B (NF-kappaB or NF-κB) pathway impacts clock function. NF-κB transcription factors play critical roles in inflammation and immunity, as well as cell proliferation, differentiation, and survival (Oeckinghaus and Ghosh, 2009; Zhang et al., 2017). NF-κB consists of a family of five related transcription factors: RELA (p65), RELB, c-Rel, NFKB1 (p105/p50), and NFKB2 (p100/p52). RELA and RELB each contain a transactivation domain and form dimers with p50 and p52, respectively, with the RELA/p50 complex representing the canonical pathway.

Proinflammatory stimuli such as tumor necrosis factor alpha (TNFα), interleukin 1 (IL-1) and bacterial lipopolysaccharides (LPS) induce the canonical NF-κB pathway. Upon stimulation, the cytosolic RELA/p50 complex translocates to the nucleus to mount a rapid response through transcriptional induction of target genes (Oeckinghaus and Ghosh, 2009; Wu and Shi, 2013; Zhang et al., 2017). As a gatekeeper, inhibitor of NF-κB (IκBα, encoded by *NFKBIA*) serves as a repressor of NF-κB by virtue of masking its nuclear localization signal and preventing its nuclear entry. IκBα is regulated by the IκB kinase complex (IKK1, IKK2, and NEMO) that phosphorylates and targets IκBα for proteasomal degradation, thereby freeing RELA/p50 to enter the nucleus. In the non-canonical RELB/p52 pathway, the p100 is phosphorylated by IKK1 and then converted to p52; the RELB/p52 dimer subsequently translocates to the nucleus (Oeckinghaus and Ghosh, 2009; Zhang et al., 2017). In this manner, RELA and RELB use different mechanisms to regulate distinct physiological functions.

Here, we characterized the effect of the canonical RELA pathway on the circadian clock. Using genetic and pharmacological approaches, we show that RELA activation altered the expression patterns of core clock genes, resulting in altered circadian rhythms in cellular models and the SCN clock, as well as circadian locomotor behavior. RELA potently repressed E-box-mediated BMAL1/CLOCK transcriptional activity at a steady-state level. Biochemical and biophysical assays revealed a direct interaction between the REL homology domain (RHD) of RELA and the C-terminal regulatory domain (CRD) of BMAL1, centering on the transactivation domain (TAD). Our data suggest that NF-κB competes with CRY1 and coactivator CBP/p300 for binding to the BMAL1 TAD. Together, these results support a significant role for NF-κB in modulating circadian clock function. Given that inflammation and innate immunity are under the control of the circadian clock (Labrecque and Cermakian, 2015; Man et al., 2016), our findings highlight the mutual regulation between these two pathways.

## RESULTS

### NF-κB alters circadian rhythms in a cellular clock model

Our functional genomic screen identified the NF-κB pathway as a clock modifier in U2OS cells (Zhang et al., 2009). For example, knockdown of *NFKBIA* (inhibitor of NF-κB, aka *IκBα*) and *IKBKB (IκBα* kinase beta, aka *IKK-β* or *IKK2*, which activates NF-κB) in these cells caused short and long periods, respectively (**Fig. S1A**). Consistent with the effect *of IκBα* knockdown and resultant NF-κB pathway activation, overexpression of a constitutively active *IKK2* mutant, *IKK2*-S177/181E (or *IKK2^CA^*) (Mercurio et al., 1997), resulted in an even shorter period length, compared to WT *IKK2* (**Fig. S1B**). Consistent with the *IKK2* knockdown effect, continuous exposure to the IKK2 inhibitor TPCA-1 (Du et al., 2012) (**Fig. 1A**), the naturally-occurring steroidal lactone, withaferin A (WA) (Kaileh et al., 2007) (**Fig. S1C**), or the orally-bioavailable NF-κB inhibitor, CAT-1041 (Hammers et al., 2016) (**Fig. S1D**) also caused longer periods in these cells, compared to DMSO control. Taken together, our genetic and pharmacological data support a role for NF-κB in regulating cell-autonomous circadian clocks.

**Figure 1.**
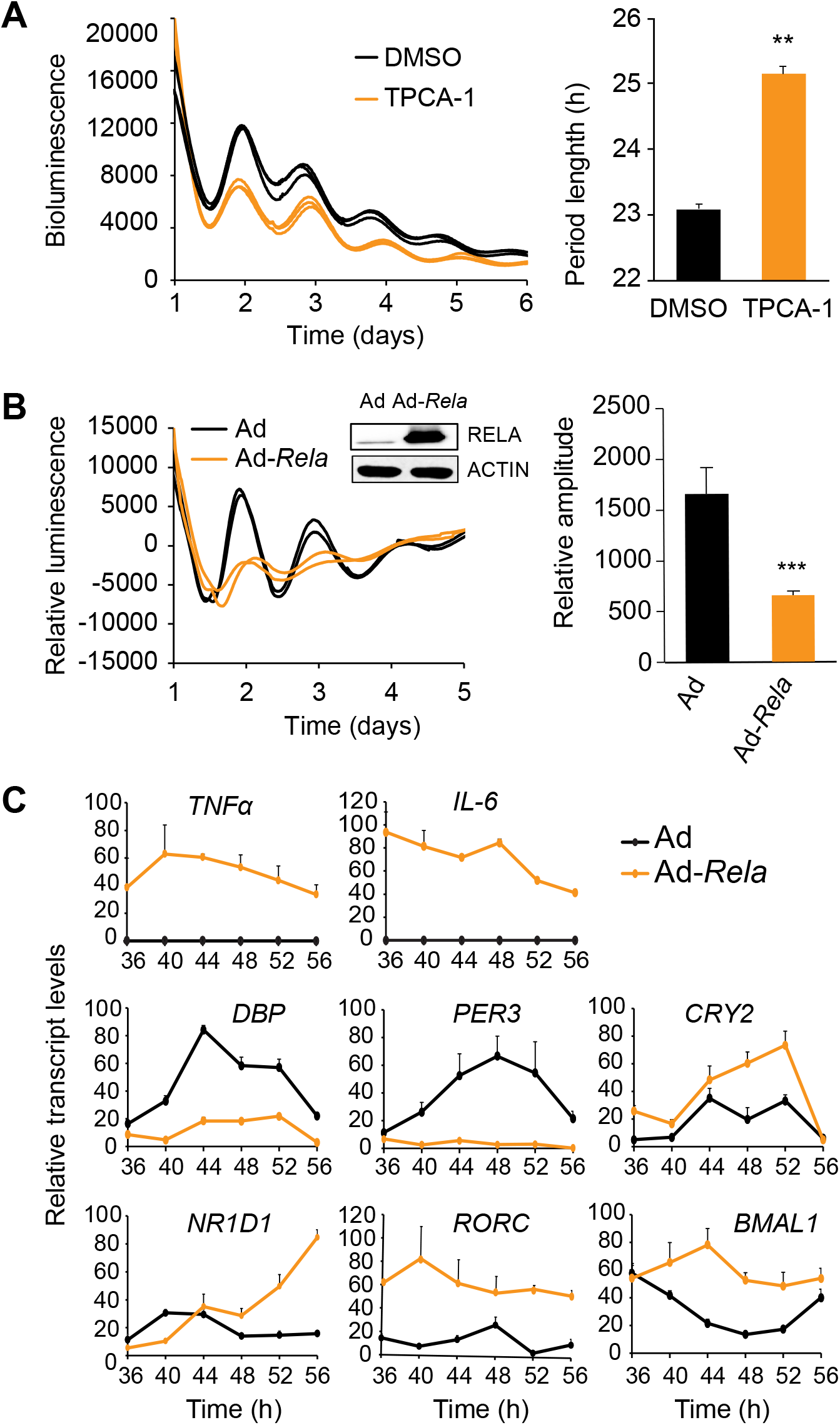
NF-κB affects clock gene expression and circadian rhythms in U2OS cells. **(A-B)** Representative circadian bioluminescence rhythms in U2OS cells harboring the *Per2-dLuc* reporter. Bioluminescence was recorded in a Lumicycle luminometer on 35-mm culture dishes and the bioluminescence data are representative of at least three independent experiments. In (A), cells were treated with IKK2-specific inhibitor TPCA-1 (5 μM). IKK2 inhibition caused long period length compared to DMSO control. Period lengths are mean ± standard deviation (SD). n = 6 independent dishes for each condition, p<0.01. In (B), cells were transduced with an adenoviral *Rela* expression vector. Ad, adenoviral vector control; Ad-*Rela*, RELA adenoviral expression vector. RELA overexpression was determined by Western blot analysis (insert). RELA overexpression reduced rhythm amplitude. Rhythm amplitudes are mean ± standard deviation (SD). n = 6 independent dishes for each condition. ** p<0.01. *** p<0.005. **(C)** RELA overexpression alters clock gene expression in U2OS cells as in (B). Cells were synchronized by dexamethasone treatment (CT0) and collected 36 hours later at CT36 at a 4 h interval for a complete cycle. Transcript levels were determined by Q-PCR and normalized to *GAPDH*. Error bars represent SD of expression levels from 3 independent wells. TNFα and IL-6 are classic targets of NF-κB and other genes are representatives of core clock components.

To more directly assess the effect of NF-κB on the clock function, we tested whether overexpression of the canonical NF-κB subunit RELA affects circadian oscillations. For this, we used the adenoviral system to overexpress the mouse *Rela* gene in U2OS cells. Compared to cells transduced with the empty vector (Ad), cells expressing Ad-*Rela* were only transiently rhythmic during the first few cycles, with a short period length, and displayed significantly decreased amplitude (**Fig. 1B**). These results suggest that NF-κB affects clock function.

### NF-κB alters core clock gene expression in U2OS cells

To determine how RELA affected circadian oscillations, we measured the expression patterns of core clock genes by quantitative PCR. We focused on the time window of 1.5-2.5 days (or 36-60 h) post synchronization when the clock phenotype was apparent but still maintained rhythmicity. As expected, *IL-6* and *TNFα*, the two classic NF-κB targets, were significantly upregulated in Ad-*Rela* cells, compared to Ad control (**Fig. 1C**), suggesting that the exogenous RELA is functional in these cells. The expression patterns of most of the core clock genes in these cells were altered (**Figs. 1C** and **S1E**). RELA drastically reduced *DBP* and *PER3* expression at all circadian times. This result is consistent with previous reports showing that LPS and TNFα, known to induce NF-κB activation, caused low levels of *DBP* in the liver and lung (Cavadini et al., 2007; Haspel et al., 2014; Hong et al., 2018). *DBP* and *PER3* are known to be regulated by the E-box (Baggs et al., 2009; Liu et al., 2008; Takahashi, 2017; Ueda et al., 2005). Further, *PER3* is also regulated by the D-box and the blunted DBP (D-box activator) likely contributed to the low *PER3* expression. In contrast, *BMAL1* was dramatically elevated at all circadian times (**Fig. 1C)**. *BMAL1* is known to be controlled by the RORE cis-element and its expression in Ad-*Rela* cells may be explained by high levels of the RORE activator *RORC* and low levels of the RORE repressor *NR1D1:* while *NR1D1* was down-regulated during peak hours (36-40 h), its expression was abnormally high during trough hours (52-56 h); in contrast, *RORC* and *RORA* levels were abnormally high throughout the circadian cycle (**Figs. 1C** and **S1E**). These expression patterns are consistent with previously reported cytokine effects (Cavadini et al., 2007; Haspel et al., 2014; Hong et al., 2018) and with the network features of the molecular clockwork (Baggs et al., 2009; Liu et al., 2008; Ueda et al., 2005).

### NF-κB alters circadian rhythms in the SCN clock and locomotor behavior

The NF-κB effect on cell-autonomous clocks raised the possibility that it also affects the master SCN clock. Indeed, we show that treatment with TPCA-1 in SCN explants derived from PER2::LUC fusion (*Per2^Luc^*) knockin mice (Yoo et al., 2004) caused long period length (**Fig. 2A**). To further test this, we infected SCN slices with Ad-*Rela*. Compared to controls that showed slightly reduced amplitude, SCN slices transduced with Ad-*Rela* displayed significantly lower amplitude and rapid loss of rhythmicity, and the period length was significantly shorter during the first few cycles of transient rhythms (**Fig. 2B**).

**Figure 2.**
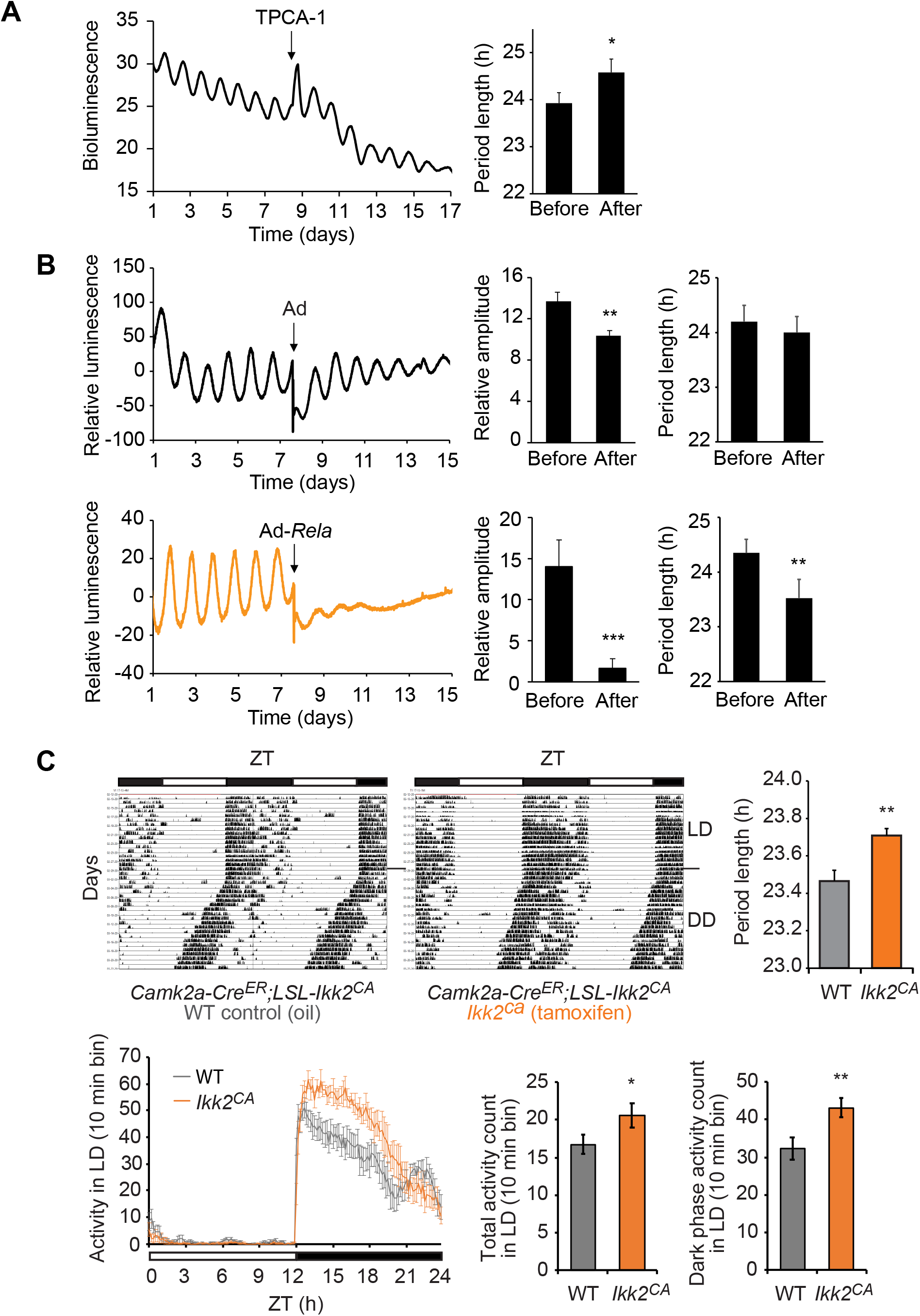
NF-κB affects circadian rhythms in the SCN and animal behavior. **(A)** Circadian bioluminescence rhythms of SCN explants from *Per2^Luc^* reporter mice treated with IKK2 specific inhibitor TPCA-1 (5 μM). Period lengths are mean ± standard deviation (SD): 23.93 h ± 0.22 before treatment; 24.58 h ± 0.29 after treatment (n = 4 SCN slices). * p<0.05. **(B)** Circadian bioluminescence rhythms of *Per2^Luc^* SCN explants expressing RELA. Ad, adenoviral vector control; Ad-*Rela*, RELA adenoviral expression vector. Compared to vector control, RELA overexpression reduced rhythm amplitude and shortened period length. Amplitude and period length data are mean ± SD (n = 4 SCN slices). ** p<0.01. *** p<0.005. **(C)** (Above) Representative double-plotted actograms of wheel-running activity rhythms in control and *Ikk2^CA^* mice. X axis: zeitgeber time (ZT) of the 12 h/12 h light/dark cycle (LD) indicated by the bar (top). Y axis: number of days during the experiment. Mice were first entrained to a regular LD cycle and then released to constant darkness (DD). Grey and white areas indicate dark and light periods. (Below) Circadian free-running parameters in DD: period lengths, total wheel-running activity, and daytime or rest phase activity. Data are mean ± SD (n = 6 for each group). * p<0.05. *** p<0.005.

Circadian animal behavior is an overt output of the SCN clock (Ko et al., 2010; Liu et al., 2007b; Mohawk et al., 2012). The NF-κB effect on the SCN clock raises the possibility that NF-κB affects the circadian locomotor activity. To test this, we obtained the *Camk2a-Cre^ER^;LSL-Ikk2^CA^* mouse line. In the *LSL-Ikk2^CA^* line, the LoxP-Stop-LoxP (LSL) cassette prevents expression of the constitutively active *Ikk2*-S177E/S181E (*Ikk2^CA^*) (Sasaki et al., 2006). In the *Camk2a-Cre^ER^* line, Cre^ER^ is conditionally expressed in neural tissues including forebrain, the hypothalamus and the SCN (Izumo et al., 2014; Madisen et al., 2010). Tamoxifen treatment induced deletion of the floxed LSL stop cassette and consequently conditional expression of IKK2^CA^ (**Fig. 2C**), leading to constitutive NF-κB activation (Sasaki et al., 2006). We performed the wheel-running locomotor activity assay to assess circadian behavioral phenotypes. Control and *Ikk2^CA^* mice were entrained to the standard 12h/12h light/dark (LD) cycle for 2 weeks, followed by release to constant darkness (DD) to monitor their endogenous locomotor activity. Like the control mice, *Ikk2^CA^* mice were able to entrain to the LD cycle (**Fig 2C**). Under the LD condition, *Ikk2^CA^* mice displayed significantly higher total activity (WT control: 16.72 ± 3.14, n = 6; *Ikk2^CA^*: 20.61 ± 4.02, n = 6; p = 0.0914) and dark phase activity than WT controls (WT control: 32.29 ± 7.17, n = 6; *Ikk2^CA^*: 43.17 ± 6.43, n = 6; p = 0.0279) (**Fig. 2C**), indicative of an altered diurnal locomotor behavior. Under the DD condition, the circadian period in *Ikk2^CA^* mice was longer than in the controls (**Fig 2C**; WT control: 23.47 h ± 0.06, n = 6; *Ikk2^CA^*: 23.71 h ± 0.04, n = 6; p = 0.0076), indicative of an altered SCN clock function. The period phenotype is modest but significant as observed in other mutant models such as *Clock^−/−^, Nr1d1^−/−^*, and *Chrono^−/−^* mice (Anafi et al., 2014; Debruyne et al., 2006; Preitner et al., 2002).

### NF-κB represses BMAL1/CLOCK transcriptional activity

The dramatic reduction of E-box genes such as *DBP* in cells overexpressing RELA raises the possibility that NF-κB more directly affects E-box transcription. To determine the RELA effect on E-box transcription, we performed steady-state reporter assays in transiently transfected HEK 293T cells. We used the *Per2::Luc* and *3xE-box::Luc* reporters, in which Luciferase (*Luc*) expression is driven either by the mouse *Per2* promoter that contains the E-boxes or by three tandem E-box repeats, respectively (Khan et al., 2012; Ramanathan et al., 2012; Yoo et al., 2005). As expected, BMAL1 and CLOCK activated E-box transcription and CRY1 effectively repressed it in both reporters (**Fig. 3A**). However, both RELA and RELB effectively repressed BMAL1/CLOCK activity, with a potency that is comparable to CRY1 (**Fig. 3A**). RELB was previously shown to repress E-box expression (Bellet et al., 2012) and our results show that RELA displayed higher E-box repression than RELB. It is noted that, RELA and RELB, known as transcriptional activators, did not activate E-box transcription (**Fig. S2A**).

**Figure 3.**
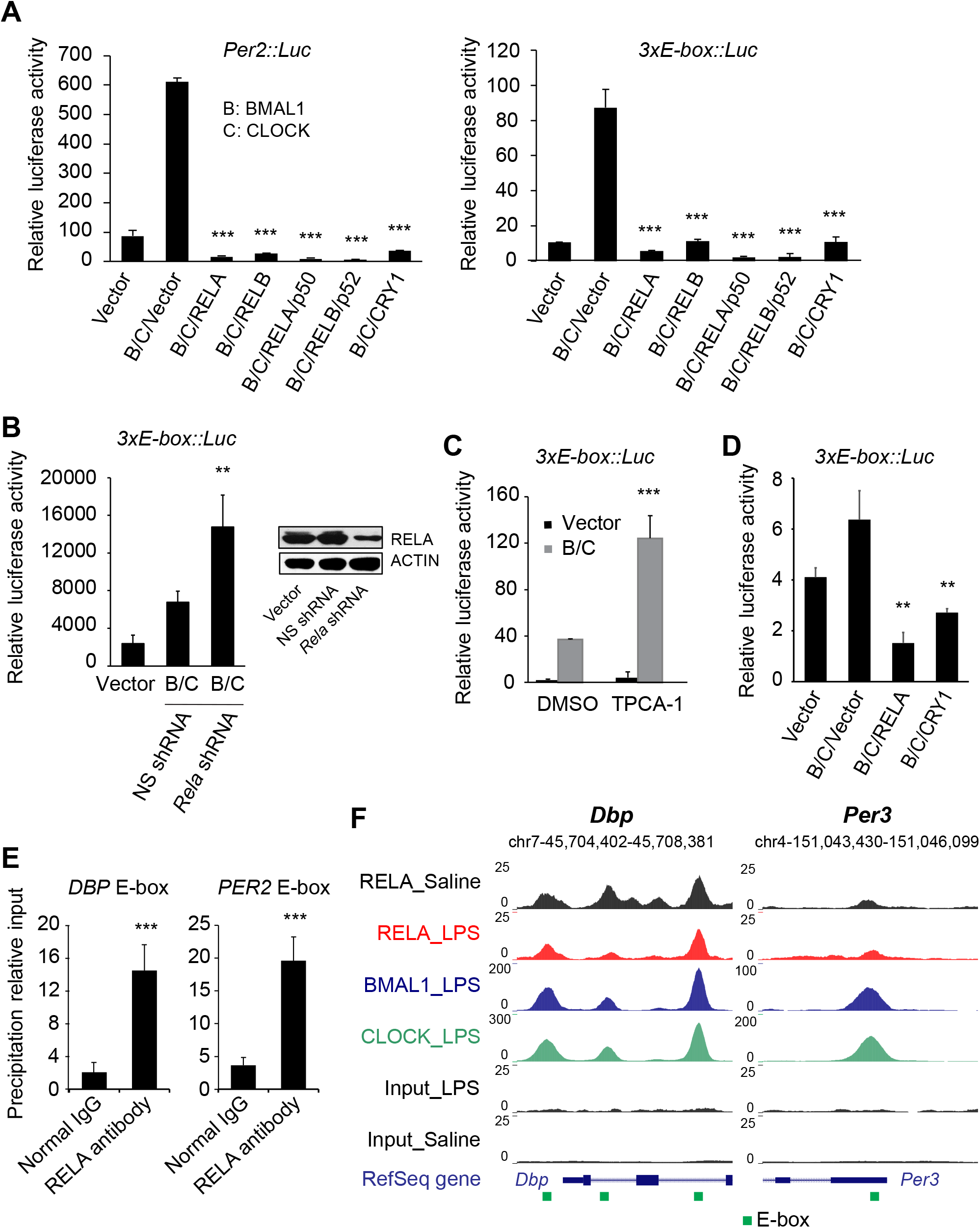
NF-κB represses E-box mediated transcriptional activity of BMAL1/CLOCK. **(A)** Steady-state luciferase reporter assay in transiently transfected 293T cells. In the *3xE-box::Luc* and *Per2::Luc* reporters, Luc expression is controlled by tandem E-box repeats or by the mouse *Per2* gene promoter containing the E-box, respectively. RELA and RELB repressed the transcriptional activity of BMAL1/CLOCK at the *Per2::Luc* (left) and *3xE-box::Luc* (right) reporters. No NF-κB-RE is present in the *3xE-box::Luc* reporter. **(B-C)** RNAi knockdown of endogenous *RELA* (B) or inhibition of IKK2 by TPCA-1 (20 μM) (C) relieved the repression of BMAL1/CLOCK by RELA. **(D)** Steady-state luciferase reporter assay in transiently transfected *Cry1^−/−^;Cry2^−/−^* fibroblasts using the *3xE-box::Luc* reporter. RELA was able to repress E-box transcription in the absence of the endogenous CRY1. **(E)** RELA associates with the E-box cis-element of clock genes *PER2* and *DBP* in U2OS cells. Chromatin immunoprecipitation (ChIP) and Q-PCR to detect binding of endogenous RELA to the promoters of *PER2* and *DBP* at the E-box. ChIP was performed with an anti-RELA or normal IgG antibody. Representative data from three independent experiments are shown. Data are mean ± standard deviation (n = 3). **(F)** Representative UCSC genome browser images of RELA, BMAL1, and CLOCK ChIP-seq tracks at the *Dbp* and *Per3* genes. Normalized tag counts are indicated on the Y-axis. Green square, E-box. ChIP-seq analysis revealed overlapping binding of BMAL1, CLOCK and RELA at the E-boxes of *Dbp* and *Per3*. Motif analysis did not find NF-κB-RE in these binding peaks, suggesting indirect binding of NF-κB to the E-box. * p<0.05. ** p<0.01. *** p<0.001.

There is a considerable level of endogenous NF-κB in 293T cells. We reason that this basal activity would repress E-box transcription. In support of this notion, we show that shRNA knockdown of endogenous *RELA* relieved this repression, leading to significantly higher levels of reporter activity (**Fig. 3B**). Similar to RNAi knockdown, inhibition of endogenous IKK2 with TPCA-1 also relieved its repression of BMAL1/CLOCK activity (**Fig. 3C**). Other inhibitors of the NF-κB pathway such as GSK143, Bay11-7082 and CAT-1041 had similar effects (**Fig. S2B**).

CRY1 and CRY2 are the canonical repressors of BMAL1/CLOCK. It is possible that NF-κB’s E-box repression is dependent on CRY1 or CRY2. To test this, we performed the reporter assay in *Cry1^−/−^;Cry2^−/−^* mouse fibroblasts (Khan et al., 2012). In these *Cry*-deficient cells, the endogenous BMAL1 and CLOCK activated the *3xE-box::Luc* reporter, which was further upregulated by cotransfected BMAL1 and CLOCK and suppressed by RELA (**Fig. 3D**). These results indicate that NF-κB represses E-box transcription in a CRY-independent manner.

### NF-κB regulates clock genes at the BMAL1/CLOCK-E-box loci

Our data support a model that NF-κB is recruited to the E-boxes of clock genes to repress BMAL1/CLOCK transcription. To test this prediction, we performed chromatin immunoprecipitation (ChIP) and quantitative PCR to determine the action of the endogenous RELA on clock genes including *PER2* and *DBP* that have validated E-boxes (Koike et al., 2012; Ueda et al., 2005). We show that, compared to IgG control, RELA displayed enriched binding to the E-boxes in the promoters of *DBP* and *PER2* (**Fig. 3E**).

Our data are consistent with a recent study showing that NF-κB targeted *Per2* and *Dbp* in the mouse liver upon LPS treatment (Hong et al., 2018). Given the rich ChIP-seq data in the study (Hong et al., 2018), we mined the publicly accessible datasets to further define how NF-κB coordinates with BMAL1/CLOCK to regulate the E-box genes. ChIP-seq analysis revealed different gene cohorts targeted exclusively by RELA or by BMAL1/CLOCK, but not both. For example, the canonical NF-κB targets such as *Nfkb1, Nfkb2, Nfkbia*, and *Rela*, displayed significant RELA binding (**Fig. S3A-B and Table S1**). Importantly, these genes were not targeted by BMAL1 or CLOCK and no consensus E-boxes were found in the regulatory regions. Similarly, other genes such as *Ccne1, Gmfb, Atxn3* and *Aven*, contain E-boxes but not NF-κB-RE, and these sites were bound by BMAL1 and CLOCK, not by RELA.

Among the peaks bound by BMAL1 (6,114), CLOCK (22,247) and RELA (13,740), 1647 showed binding by all three factors, representing triple overlapping binding. All the core clock genes that are known to be regulated by BMAL1 and CLOCK via the E-box are represented in this cohort, including *Per1, Per2, Per3, Cry1, Cry2, Dbp, Nr1d1/Rev-erbα, Nr1d2/Rev-erbβ, Rorc/Rorγ*, and *Ciart/Chrono* (**Fig. 3F, Fig. S3C-D and Table S1**), as well as *Tef, Nfil3/E4bp4, Bhlhe40/Dec1* and *Bhlhe41/Dec2* (**Table S1**). Among these genes, *Per1* and *Per2* peaks harbor both E-box and NF-κB-RE (**Fig. S3C and Table S1**), suggesting that their transcription may be regulated by RELA either directly through binding to NF-κB-RE or indirectly through binding to the BMAL1/CLOCK complex. Importantly, however, the triple binding peaks in other clock genes contain E-boxes only, but no consensus NF-κB-REs, indicative of indirect RELA binding on these chromatin sites. These data strongly support our model that NF-κB indirectly represses E-box transcription through complex formation with BMAL1/CLOCK at the E-box.

### NF-κB interacts with BMAL1 in the BMAL1/CLOCK complex

Given that many genes (e.g. *TNFα, IL-6, BMAL1*) are upregulated by RELA (**Fig. 1C**), NF-κB’s repression on the E-box is unlikely caused by general transcriptional repression. Although known as an activator in the inflammatory response, RELA can repress transcription of some target genes in an HDAC-dependent manner (Perkins, 2007). However, inhibition of HDACs with chemical inhibitors such as nicotinamide and valproic acid did not attenuate RELA repression of E-box transcription (**Fig. S2C**), arguing against an HDAC-dependent mechanism. Previous studies detected interactions between RELA and CLOCK at NF-κB target genes (Spengler et al., 2012), and between RELB and the BMAL1/CLOCK complex (Bellet et al., 2012). Our data also show that RELA associated with the E-boxes of clock genes (**Fig. 3E**). As the *3xE-box::Luc* reporter does not contain NF-κB cis-response elements and alone is not responsive to NF-κB, it is likely that NF-κB affects E-box transcription via an indirect mechanism. In light of these observations, we hypothesize that RELA repression occurs through direct interactions with BMAL1 and/or CLOCK.

We show that Myc-RELA interacts with Flag-BMAL1 by co-immunoprecipitation (co-IP) and Western Blot analysis (**Fig. 4A**). The RELA and CLOCK interaction was weak but appeared stronger in the presence of BMAL1 (**Fig. S4A-C**). Co-IP detected an endogenous interaction between BMAL1 and RELA in WT and *Clock^−/−^* cells, but not in *Bmal1^−/−^* cells (**Figs. 4B** and **S4D**). These results suggest that the complex formation is dependent on BMAL1, but not CLOCK, and the RELA and CLOCK interaction occurred indirectly through BMAL1.

**Figure 4.**
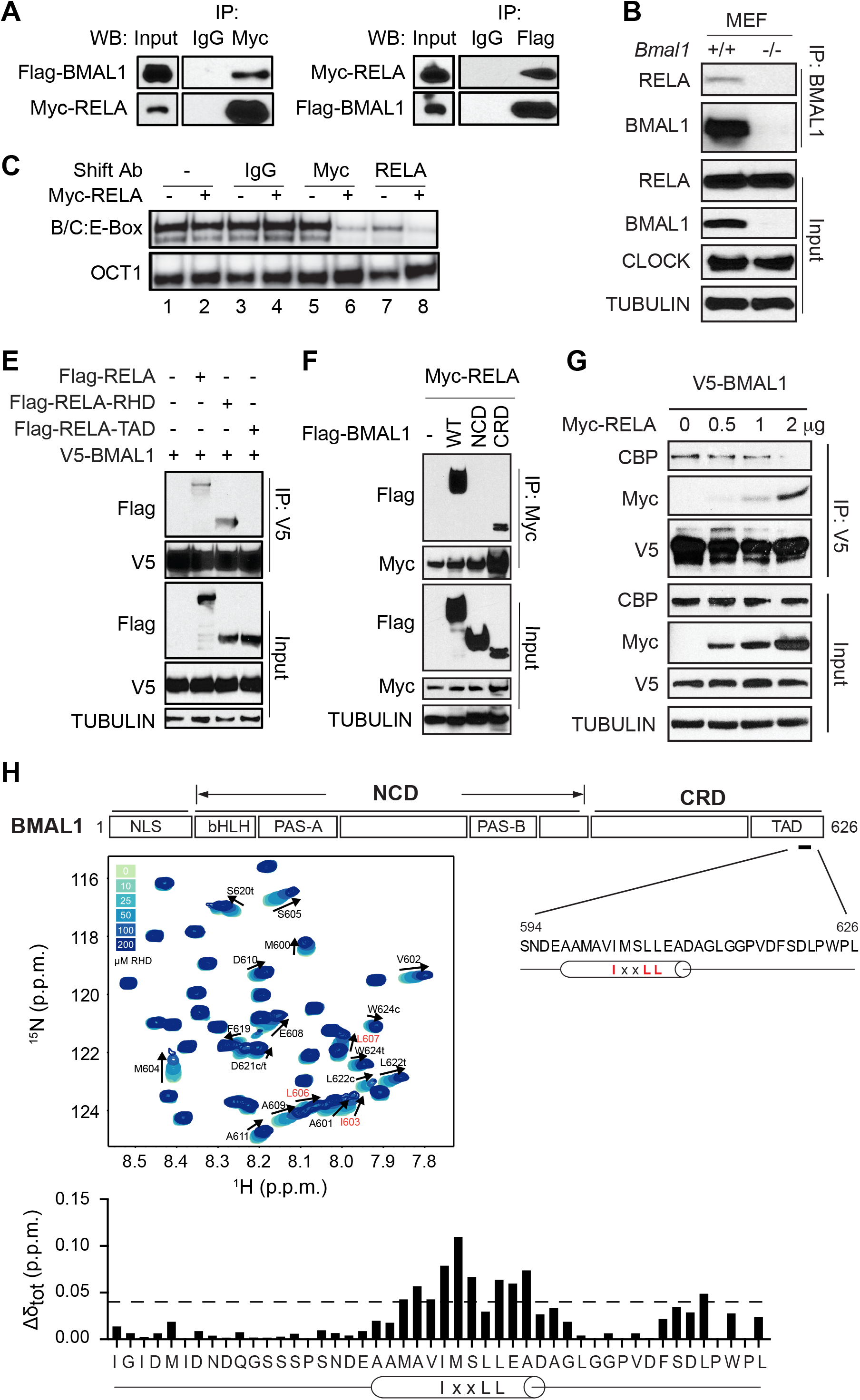
The RELA RHD interacts with the C-terminal transactivation domain of BMAL1. **(A-B)** Co-immunoprecipitation assays detect the interaction between RELA and BMAL1 in co-transfected 293T cells (A) and between the endogenous proteins in fibroblasts (B). The intrinsic interaction between RELA and BMAL1 is independent on CLOCK. **(C)** RELA associates with the BMAL1/CLOCK:E-box complex (C), indicative of a larger complex formation which caused a supershift. Oct1 oligonucleotide was used as control. The complex was reduced by the anti-Myc antibody, likely caused by a supershift that was not revealed in the gel. **(E-F)** Co-immunoprecipitation and Western blot analysis to detect the interaction between the RELA RHD and the BMAL1 CRD. 293T cells were co-transfected with tagged BMAL1 and RELA proteins as indicated. **(G)** Co-immunoprecipitation and Western blot analysis to show competition for interaction with BMAL1 between CBP and RELA. **(H)** Backbone chemical shift perturbations (Δδ_tot_) of ^15^N-BMAL1 C-terminal TAD with stoichiometric RELA RHD. Top panel: Schematic diagram of domain structure of BMAL1. The N-terminal core domain (NCD) contains the bHLH, PAS-A and PAS-B domains. The C-terminal regulatory domain (CRD) contains a transactivation domain (TAD) near the C-terminus featuring the highly conserved IxxLL helical motif. Residues within the IxxLL helical motif showed chemical shifts, indicative of interaction with RELA. Middle: Central region of ^15^N-^1^H HSQC spectra showing titration of 100 μM ^15^N-BMAL1 TAD in the presence of increasing concentrations of the RELA RHD (light to dark blue; solid arrow). Peak assignments are indicated for residues in the BMAL1 TAD that undergo chemical shift perturbation upon the addition of RELA RHD. Bottom: Quantification of backbone chemical shift perturbations (Δδ_TOT_) of ^15^N-BMAL1 TAD from the titration point with 200 μM RELA RHD. Dashed line, Δδ_TOT_ significance cutoff of 0.04 p.p.m. (parts per million).

Using electrophoretic gel mobility shift assay (EMSA), we show that both endogenous and ectopically expressed BMAL1 and CLOCK formed a complex with a radio-labeled 14 bp E-box duplex probe (**Figs. 4C** and **S4E**). As shown, this ternary complex at the E-box required both BMAL1 and CLOCK, as the complex was absent in mouse fibroblasts deficient in *Bmal1* or *Clock* (**Fig. S4E**). This binding was effectively competed out with 100-fold excess of unlabeled (or cold) wild type E-box oligomer, but not with cold mutated E-box, confirming the specificity of the E-box complex formation (**Fig. S4E**). The complex in the EMSA did not show a noticeable difference with or without Myc-RELA, suggesting that RELA itself does not disrupt the BMAL1/CLOCK:E-box complex (**Fig. 4C**, lanes 1-2). An anti-Myc antibody, but not normal IgG, significantly decreased the complex signal, suggesting that Myc-RELA is part of the BMAL1/CLOCK:E-box complex (**Fig. 4C**, compare lane 4 with lane 6). The complex was reduced by the anti-Myc antibody, likely caused by a supershift that was not revealed in the gel. Indeed, anti-RELA antibody caused similar changes for both endogenous and exogenous RELA (**Fig. 4C**, compare lanes 4 and 8, and lanes 3 and 7). Taken together, these data support a model in which NF-κB binds directly to the BMAL1/CLOCK:E-box complex to repress E-box transcription rather than inducing disassembly of the complex.

### The RELA RHD domain interacts with the BMAL1 C-terminal regulatory domain

NF-κB functions primarily as a transcriptional activator (Perkins, 2007). We asked whether the NF-κB effect on the E-box is dependent on its transcriptional activity. NF-κB contains an N-terminal Rel homology domain (RHD) and C-terminal transactivation domain (TAD) (Oeckinghaus and Ghosh, 2009; Zhang et al., 2017) (**Fig. S5A**). Mutation of Serine 281 within the RHD to glutamate (S281E) abolishes NF-κB transcriptional activity (Maier et al., 2003). Using the steady-state reporter assay, we show that RELA-S281E lost its transcriptional activity to induce the *NF-κB-RE::dLuc* reporter, but was still able to repress the *3xE-box::Luc* reporter, despite weaker repression than WT RELA (**Fig. S5B**). To determine which RELA domain underlies its repression, we generated two truncation constructs, RELA-RHD and RELA-TAD. We found that RELA-RHD, but not RELA-TAD, retains E-box repression activity in the *3xE-box::Luc* reporter assay (**Fig. S5B**). Taken together, these results suggest that the classic NF-κB transcriptional activator activity is not required for its repression of the E-box.

Since RELA and BMAL1/CLOCK form a complex on the E-box, we next determined the specific domains that underlie the interaction. Co-IP detected an interaction between BMAL1 and the RELA RHD, but not the TAD (**Fig. 4E**). Conversely, we asked which BMAL1 region associates with RELA. The N-terminal core domain (NCD) of BMAL1 contains the bHLH and PAS domains, responsible for dimerization and DNA binding (Huang et al., 2012). Our recent studies found that part of the C-terminal regulatory domain (CRD) coordinates binding to CRY1 and coactivator CBP/p300, and their dynamic interactions plays a critical role in enabling circadian oscillations (Gustafson et al., 2017; Xu et al., 2015). Intriguingly, we show that, like CRY1, RELA also interacted with the BMAL1 CRD (**Fig. 4F**). These data suggest that RELA impacts clock function likely through competitive binding with CBP/p300. To test this hypothesis, we performed co-IP and Western blot analysis and showed that V5-BMAL1 interacts with both Myc-RELA and CBP. Importantly, Myc-RELA was able to effectively compete with CBP for BMAL1 binding, with increasing amounts of Myc-RELA correlating with decreased amounts of CBP (**Fig. 4G**). These results further support the mechanism in which RELA competes with CBP for binding to the BMAL1 CRD.

### NF-κB binds to the BMAL1 C-terminal transactivation domain

In our previous study, we showed that CRY1 and CBP compete for binding to the transactivation domain (TAD) within the BMAL1 CRD and the shared binding sites center on the IxxLL helical motif within the α-helix of the TAD and on a second cluster of residues in the distal C terminus (Gustafson et al., 2017; Xu et al., 2015) (**Fig. 4H**). Our results showing that NF-κB binds to the BMAL1 TAD raised the possibility that the same IxxLL motif is involved. To determine the precise binding site of NF-κB on the BMAL1 TAD, we collected ^15^N HSQC NMR spectra of ^15^N-labeled TAD in the presence or absence of the RELA RHD and showed that titration of the RELA RHD led to dose-dependent chemical-shift perturbations at residues within the helical motif of the TAD and the distal C terminus, overlapping directly with regions that bind CRY1 and CBP (**Fig. 4H**). These results confirm the direct interaction between BMAL1 and NF-κB and demonstrate that CBP, CRY1 and NF-κB binding all converge on overlapping binding sites on the BMAL1 TAD.

## DISCUSSION

### NF-κB is a circadian clock modifier

Our genome-wide RNAi screen uncovered many genes, representing various cellular pathways and functions, that input to and modify the clock (Zhang et al., 2009). The past decade has seen several examples of the integration of circadian clocks directly with cell physiology: AMP-activated protein kinase (AMPK) (Lamia et al., 2009), mTOR and autophagy in response to nutrient and growth signals (Feeney et al., 2016; Lipton et al., 2017; Ramanathan et al., 2018; Toledo et al., 2018; Zandra E Walton et al., 2018), NAD^+^-dependent deacetylase sirtuin-1 (SIRT1) (Asher et al., 2008; Nakahata et al., 2009, 2008; Ramsey et al., 2009), oxidative stress inducible transcription factor NRF2 (Rey et al., 2016; Wible et al., 2018), and hypoxia inducible factor HIF1α (Adamovich et al., 2017; Peek et al., 2017; Wu et al., 2017). In the present study, we identified NF-κB as a clock modifier and characterized the mechanism of how NF-κB regulates circadian clock function. NF-κB appears to bind to the regulatory sequences of core clock genes to affect their expression even under non-stress conditions. We show that it represses E-box transcription through direct binding to the BMAL1 TAD. In the U2OS cellular clock model, NF-κB activation and inhibition shortened and lengthened circadian rhythm period lengths, respectively, whereas suboptimal NF-κB activity correlated with low amplitude phenotypes (**Figs. 1–2** and **S1**). NF-κB perturbation also altered the SCN clock and circadian behavioral rhythms, as seen in *Ikk2^CA^* (**Fig. 2**) and *Ikk2^−/−^* mice (Hong et al., 2018). Taken together, basal NF-κB activity under normal conditions is critical for maintenance of circadian homeostasis and increased NF-κB activity impacts the clock function.

The long period phenotype in *Ikk2^CA^* mice (NF-κB constitutive activation) is mechanistically consistent with the short period phenotype observed in *Ikk2^−/−^* mice (Hong et al., 2018). However, these behavioral phenotypes (e.g. long period in *Ikk2^CA^* mice) are surprising, as cells and SCN explants showed short period phenotype. This phenotypic difference between animal behavior and cell and tissue clocks (as in U2OS and SCN) suggest possible involvement of systemic signals such as those in the neuroendocrine and autonomic neural systems. Of relevance, there exists a reciprocal regulation between innate immunity and sleep disturbance (Irwin, 2015), and constitutive NF-κB activation or inactivation may alter neuroimmunology at the organismal level, contributing to cell/tissue non-autonomous behavioral phenotypes.

### Mechanisms of E-box repression by NF-κB

NF-κB could affect clock function by several mechanisms: binding to NF-κB-REs present in clock genes to directly upregulate gene expression, recruiting HDACs to modify chromatin and repress transcription (Perkins, 2007), and interacting with BMAL1/CLOCK bound at the E-box to indirectly repress gene expression. While upregulation of canonical NF-κB targets via the NF-κB-RE correlated with increased RNA Pol II binding and increased H3K27 acetylation, NF-κB repression of BMAL1/CLOCK activity at the E-box was correlated with decreased Pol II binding and decreased H3K27 acetylation (Hong et al., 2018). It was suggested in that study that LPS affected the clock repressors such as *Per*, *Cry, E4bp4* and *Nr1d1*, not the *Bmal1, Clock, Dbp* and *Ror* activators. Importantly, however, the high-resolution ChIP-seq data revealed that most of the BMAL1/CLOCK/RELA triple binding peaks, including not only the clock repressors but also the activators such as *Dbp, Tef* and *Rorc*, contain only E-boxes, suggesting indirect binding of NF-κB at the E-box (**Figs. 3** and **S3**, **Table S1**). This is evident in the *3xE-box::Luc* reporter assay, in which the promoter sequence, unlike the endogenous genes, does not contain NF-κB-REs or other confounding factors (**Fig. 3**). Further, genes such as *Bmal1* and *Clock* that are not regulated via the E-box did not show RELA binding. Intriguingly, RELA displayed binding peaks at the E-boxes in saline controls, independent of LPS treatment. Some of these peaks were slightly shifted by LPS and even displayed decreased binding for all three factors (**Figs. 3** and **S3**) (Hong et al., 2018). These observations suggest that LPS-induced NF-κB has some remodeling effect on the BMAL1/CLOCK-E-box complex. Future studies are needed to investigate how BMAL1, CLOCK and NF-κB coordinate to regulate E-box transcription.

Compared to NF-κB transcriptional activation, its repression mechanism is much less understood (Oeckinghaus and Ghosh, 2009; Perkins, 2006). Of relevance, acetylation of RELA by CBP/p300 at specific sites can have both positive and negative effects on DNA binding and transcription, and so do phosphorylation and dephosphorylation. Interestingly, RELA can also recruit HDACs that modify chromatin to repress transcription (Perkins, 2007). However, HDAC inhibitors did not affect RELA repression of BMAL1/CLOCK **(Fig. S2C**), suggesting that HDAC activity is not required for E-box repression. Our work showed that the BMAL1 CRD coordinates interactions with CRY1 and CBP/p300 which plays a key role in enabling circadian oscillations (Gustafson et al., 2017; Xu et al., 2015). In this study, we show that, like CRY1, RELA also interacts with the BMAL1 CRD. Thus, RELA impacts clock function likely through competitive binding with CRY repressors. Recent studies have made important progresses in our mechanistic understanding of CRY repression which involves dynamic interactions with the BMAL1 CRD and CLOCK PAS domains (Fribourgh et al., 2020; Gustafson et al., 2017; Michael et al., 2017; Xu et al., 2015). As repressors of BMAL1/CLOCK at the E-box, RELA and CRY may share some aspects of the repression mechanism which warrant future studies.

### NF-κB underlies mutual regulation between circadian clocks and inflammation

Both the innate and adaptive immunological functions have prominent circadian rhythms, including gene expression, synthesis and secretion of cytokines and chemokines, and immune cell functions (Labrecque and Cermakian, 2015; Man et al., 2016). Within innate immunity, recent studies have documented prominent circadian rhythms in monocytes and macrophages (Labrecque and Cermakian, 2015). Toll-like receptors and cytokine and chemokine display circadian oscillations. Several lines of evidence support circadian regulation of NF-κB activity. First, key players in the NF-κB pathway (e.g., IκBα) are rhythmically expressed in several peripheral tissues including the liver and lung, exhibiting a circadian phase similar to *Per1* (Koike et al., 2012; Zhang et al., 2014). Second, CLOCK was shown to be recruited by RELA at the NF-κB-RE to influence NF-κB target genes (Spengler et al., 2012), adding another layer of circadian regulation of NF-κB activity. NF-κB signaling was altered in *Clock* mutant mice and in *Cry*-deficient mice (Lee and Sancar, 2011; Spengler et al., 2012). In a different twist, RELA activation was shown to be regulated through CRY repression of adenylyl cyclase and consequently inhibition of PKA-mediated RELA phosphorylation (Narasimamurthy et al., 2012). RORα, a key activator of the RORE loop, upregulates IκBα expression, thereby hindering NF-κB nuclear translocation and activity (Delerive et al., 2001). Third, the *NF-κB-RE::dLuc* reporter assay showed that NF-κB displays circadian activities in the liver (Spengler et al., 2012). These studies support mutual regulation between the circadian and the inflammatory pathways.

Several studies in recent years have demonstrated the effects of inflammation on clock function. Inflammation induced by endotoxin treatment with LPS, IL-1 or TNFα altered clock gene expression in the SCN (Cavadini et al., 2007; Okada et al., 2008), the lung (Haspel et al., 2014; Yang et al., 2014), liver (Cavadini et al., 2007; Hong et al., 2018; Okada et al., 2008), and fibroblasts (Cavadini et al., 2007), as well as locomotor activity (Hong et al., 2018; Leone et al., 2012, 2006; Marpegán et al., 2005; Paladino et al., 2014). LPS-induced lung and liver inflammation caused a reprograming of core clock genes and outputs and led to a short period length phenotype (Haspel et al., 2014; Hong et al., 2018), consistent with our data. In a mouse model of chronic obstructive pulmonary disease (COPD), cigarette smoke exposure combined with viral infection disrupts clock function and leads to enhanced inflammatory responses in the lung and reduced locomotor activity (Sundar et al., 2015). Further, endotoxin was shown to suppress clock gene expression in human peripheral blood leukocytes (Haimovich et al., 2010). At the molecular level, both RELA and RELB can directly interact with BMAL1 and repress BMAL1/CLOCK activity, thus providing a mechanistic basis for the inflammatory effect on circadian clock function.

## EXPERIMENTAL PROCEDURES

### Plasmids

The expression constructs pLV7-CAG-1xFlag-*Bmal1* (full length), 1xFlag-*Bmal1*-NCD containing the bHLH, PAS-A and PAS-B domains, 1xFlag-*Bmal1*-CRD containing the G and H domains, *Bmal1*-V5, and 3xFlag-*Clock* were cloned as described previously (Ramanathan et al., 2018; Xu et al., 2015). The plasmids encoding Flag-*Rela* (plasmid #20012), Flag-*Relb* (#20017), Flag-*p50*(#20018) and Flag-*p52* (#20019) were obtained from Addgene. *Rela*-S281E mutant was generated with primers 5’-tctgat cgcgagctcgaggagcccatggagttc-3’ and 5’-gaactccat gggctcctcgag ctcgcgatcaga-3’. The PCR product was inserted into p3xFlag-CMV-10 vector at HindIII and EcoRI sites. *Rela*-RHD was generated with primers 5’-agctaagcttatggacgatctgtttcc-3’ and 5’-agctgcggc cgcttacatgatactcttgaaggtctc-3’, *Rela*-TAD with 5’-agctaagcttgagacctt caagagtatcatg-3’ and 5’-agctgcggccgcttaggagctgatctgactcaa-3’. The amplified products were cloned into p3xFlag-CMV-10 vector at HindIII and NotI sites. pLV7-CAG-*Ikk2* WT and *Ikk2*-S177/181E were generated as previously described (Mercurio et al., 1997; Ramanathan et al., 2018). The pGL3-*Per2::Luc* and pGL3-*3xE-box::Luc* reporters used in 293T transient transfection reporter assay were described previously (Khan et al., 2012; Ukai-Tadenuma et al., 2011; Xu et al., 2015).

For adenoviral gene expression, pShuttle-CMV (Addgene #16403) and AdEasier-1 cells (#16399) were purchased from Addgene. Full length *Rela* gene was inserted into pShuttle-CMV with SalI and NotI to obtain pShuttle-CMV-*Rela*. pShuttle-CMV and pShuttle-CMV-*Rela* were linearized with PmeI, and transformed into competent AdEasier-1 bacterial cells for recombination. The recombinants are named Ad for vector control and Ad-*Rela* for *Rela*. Recombinant adenoviruses were prepared and used as described previously (Luo et al., 2007).

### Animal and wheel-running locomotor activity assay

Mice were housed in the animal facility at the University of Florida. All animal experiments were conducted according to the National Institutes of Health Guide for the Care and Use of Laboratory Animals and approved by the Institutional Animal Care and Use Committee at the University of Florida. SCN organotypic slices were dissected and cultured in explant medium as previously described (Liu et al., 2007b; Ramanathan et al., 2018).

Mice conditionally expressing constitutively active IKK2 (*Ikk2^CA^*) in neural tissues including the forebrain and SCN (Camk2a-Cre^ER^;LSL-*Ikk2^CA^* mice, abbreviated as *Ikk2^CA^* mice) were generated by crossing the LSL-*Ikk2^CA^* mice (in which a LoxP-flanked STOP cassette prevents transcription of Flag-tagged *Ikk2^CA^* and EGFP) (Sasaki et al., 2006) with the Camk2a-CreER mice (Madisen et al., 2010) (gift of Dr. Sylvain Dore, Department of Anesthesiology). Both lines were obtained from the Jackson Laboratory. Mice of ^~^3 months old were treated with either corn oil (control) or tamoxifen (75 mg per kilogram of body weight) via i.p. injection for 5 consecutive days. Mice were used for experiments 2 weeks after tamoxifen treatment. Western blot with anti-Flag antibody was performed to determine transgene expression.

Mice were individually housed in cages equipped with running wheels and locomotor activities were recorded as previously described (Liu et al., 2007b; Ramanathan et al., 2018). Briefly, mice were entrained to a standard 12hr/12hr light/dark cycle for 2 weeks days and then released to constant darkness (DD) for 2-3 weeks. Wheel-running activities were recorded and analyzed using the ClockLab program (Actimetrics).

### Cell culture, transfection and steady-state reporter assays

Cell culture and growth conditions for HEK 293T, U2OS and MMH-D3 cells were performed as previously described (Ramanathan et al., 2014; Xu et al., 2015; Zhang et al., 2009). Briefly, 293T and U2OS cells were maintained in Dulbecco’s Modified Eagle Medium (DMEM) (HyClone) supplemented with 10% fetal bovine serum. MMH-D3 cells were grown in RPMI Medium1640 (HyClone) supplemented with 10% fetal bovine serum, 16 ng/ml insulin like growth factor-II (IGF-II), 55 ng/ml epidermal growth factor (EGF), and 10 μg/ml Insulin, as previously described (Ramanathan et al., 2014; Ramanathan and Liu, 2018). For the reporter assay, 293T cells were cultured and seeded in 96-well plates, and transfected with desired plasmids using Lipofectamine 2000 (Invitrogen). Cells were harvested 24 h later and assayed with the Dual-Glo Luciferase Assay System (Promega). Firefly luciferase activity was normalized to Renilla luciferase as an internal control for transfection efficiency, as detailed in our previous studies (Khan et al., 2012; Sato et al., 2006; Xu et al., 2015).

### Bioluminescence recording and data analysis

We used a LumiCycle luminometer (Actimetrics) for luminescence recording as previously described (Ramanathan et al., 2018, 2012; Wible et al., 2018). Briefly, cells were grown to confluence in 35-mm dishes and synchronized with 200 nM dexamethasone before bioluminescence recording. For adenoviral infection, the cells were first synchronized with dexamethasome and then infected with adenovirus for 4 h, followed by the bioluminescence rhythm assay. The recording medium contained 1x DMEM, 10 mM HEPES, 144 mM NaHCO_3_, 1% fetal bovine serum, 1x B-27 and 0.5 mM luciferin, buffered to pH 7.4. Three independent dishes/samples were used for each condition. Raw luminescence data (counts/s) as a function of time (days) were analyzed with the LumiCycle Analysis program (version 2.53, Actimetrics) to determine circadian parameters. Briefly, raw data were fitted to a linear baseline, and the baseline-subtracted data were fitted to a sine wave (damped), from which period length and goodness-of-fit and damping constant were determined. For samples that showed persistent rhythms, goodness-of-fit of 90% was usually achieved. For amplitude analysis, raw data from day 3 to day 5 were fitted to a linear baseline, and the baseline-subtracted (polynomial number = 1) data were fitted to a sine wave, from which amplitude values were obtained.

### Quantitative PCR analysis

For quantitative PCR (qPCR) analysis, U2OS cells were synchronized using 200 nM dexamethasone for 2 h and infected with adenovirus expressing Ad vector control or Ad-*Rela*. 4 h later, cells were changed to regular DMEM culture medium. After 36 h, cells were harvested at 4 h intervals for a complete circadian cycle. RNA extraction, reverse transcription, and quantitative real-time PCR were performed as previously described (Ramanathan et al., 2018, 2014; Wible et al., 2018). SYBR Green PCR master mix (Thermo Scientific) was used in qPCR. The primers used in qPCR analysis are listed in **Table S2**. Transcript levels for each gene were normalized to *GAPDH* and values were expressed as percentage of expression in control cells.

### Electrophoretic mobility shift assay

Electrophoretic mobility super-shift assay (EMSA) was performed as done previously (Wang et al., 2015). In brief, 20 μg of total cell extract was incubated with ^32^P-labeled double-stranded oligonucleotides containing an E-box (CGCGCAAAGC<u>CATGTG</u>CTTCCCCCT) at room temperature for 20 min. The samples were resolved on acrylamide gel electrophoresis and quantified with a Cyclone phosphoimager (Perkin Elmer). For supershift assay, cell extracts were incubated with 1 μg of antibody in 10 μl of EMSA reaction buffer on ice for 30 min, followed by procedures for EMSA. Oct1 oligonucleotide (Promega) was used as control. Wild type and mutant E-box sequences used as cold competition probes are listed in **Table S3**.

### Immunoprecipitation and immunoblotting

Cell lysate preparation and immunoblotting were performed as previously described (Lee, 2007a, 2007b; Ramanathan et al., 2018; Wang et al., 2015) with minor modifications. Briefly, cells were harvested by trypsinization and immediately lysed in RIPA lysis buffer containing cocktails of proteases inhibitors (Roche) and phosphatase inhibitors (Sigma). The primary antibodies used in this experiment are as following: guinea pig antibodies against BMAL1 and CLOCK from Choogon Lee’s lab; rabbit antibodies against RELA (Cell Signaling), Myc (Cell Signaling), Flag (Sigma), HA (Sigma); and Actin and Tubulin were from Santa Cruz Biotech.

Immunoprecipitation was performed as previously described (Lee, 2007c) with minor modifications. In brief, cells were lysed in 10% PBS and 90% IP lysis buffer (20 mM Tris, pH 7.0, 250 mM NaCl, 3 mM EDTA, 3 mM EGTA, 0.5% Nonidet P-40, 2 mM DTT, 0.5 mM PMSF, 20 mM β-glycerol phosphate, 1 mM sodium orthovanadate, 1 μg/ml leupeptin, 1 μg/ml aprotinin, 10 mM *p*-nitrophenyl phosphate, and 10 mM sodium fluoride). 5% of total lysate was used as input control samples. The rest of the lysate was incubated with 1 μg of primary antibody on a rotator at 4°C overnight. Next day, protein G-sepharose was added to the mixture and incubated at 4°C for 4 hr. Finally, protein G-sepharose enriched complexes were resolved on SDS-PAGE and analyzed by Western blot analysis.

Chromatin immunoprecipitation (ChIP) was performed as previously described (Lee, 2007c; Wang et al., 2015) with minor modifications. U2OS cells were synchronized using 200 nM dexamethasone for 2 hours and then infected by adenovirus of Ad vector control or Ad-*Rela*. 4 hours later, cells were changed to regular DMEM culture medium. After 36 hours, cells were fixed for 10 min at room temperature in 1x PBS containing 0.5% formaldehyde and quenched with 0.125 M glycine for 15 min. After washing with cold PBS, the cells were scraped into 1x PBS in a 15 mL conical tube and pelleted at 1,500 rpm for 10 min at 4°C. The pellet was suspended and lysed in lysis buffer (5 mM PIPES, pH 8.0, 85 mM KCl, 0.5% Nonidet P-40) for 10 min on ice. Cell lysates were sonicated with eight 20-second pulses with 20-second pauses. Cell nuclei were pelleted at 1,300 rpm for 10 min at 4°C and resuspended in 1 mL nuclei lysis buffer (50 mM Tris-Cl, pH 8.0, 10 mM EDTA, 1% SDS, supplemented with protease inhibitors), followed by incubation on ice for 10 min and then sonication with six 15-second pulses with 45-second pauses. The samples were centrifuged at 15,000 rpm at 4°C for 5 min and supernatants were used for ChIP. Primers used in qPCR to amplify the E-box region of *PER2* and *DBP* genes were designed based on previous studies (Ueda et al., 2005) and are listed in **Table S4**.

### NMR spectroscopy

All proteins were expressed in *Escherichia coli* Rosetta2 (DE3) cells from a pET22b vector backbone. Recombinant proteins for RELA-RHD (1-318) and BMAL1 TAD (579-626) were cloned from the mouse *Rela* or *Bmal1* genes, and each had an N-terminal TEV-cleavable His-GST tag. Cells were grown in Luria Broth media (for natural abundance RELA-RHD) or M9 minimal media with ^15^N-ammonium chloride for uniform incorporation of the stable isotope for NMR of BMAL1 TAD. All cultures were grown at 37°C until an O.D._600_ of ^~^0.8 was reached, after which expression was induced with addition of 0.5mM IPTG. Cultures were grown at 18°C for an additional 16-20 hours. Cell pellets for the BMAL1 TAD were lysed in buffer containing 50 mM Tris, pH 7.5, 300 mM NaCl and 20 mM imidazole for affinity purification using Ni-NTA resin (Qiagen) as we have done previously (Xu et al., 2015). Imidazole-eluted protein was buffer exchanged to a low-imidazole buffer by desalting column or diafiltration with an Amicon stirred-cell concentrator under nitrogen pressure. The His-GST tag was removed via TEV protease overnight at 4°C. Cleaved protein was separated from TEV and the His-GST tag with a Ni-NTA column and proteins were further purified by size-exclusion chromatography on Superdex 75 16/600 (GE Life Sciences) in NMR buffer (10 mM MES, pH 6.5, and 50 mM NaCl). For RELA-RHD, cells were lysed using a high-pressure extruder (Avestin) in buffer containing 50 mM Tris, pH 7.5, 300 mM NaCl, 1mM EDTA, 5 mM BME. Affinity purification was carried out with Glutathione Sepharose 4B resin (GE Healthcare). The His-GST tag was removed via TEV protease overnight at 4°C and cleaved protein was separated from the TEV and the His-GST tag with a Ni-NTA column. Protein was further purified with size-exclusion chromatography on HiLoad 16/600 Superdex 75 prep grade column (GE Healthcare) in NMR buffer. Proteins were aliquoted and frozen in liquid nitrogen for long-term storage at −80°C.

NMR experiments were conducted on a Varian INOVA 600-MHz spectrometer at 25 °C equipped with ^1^H, ^13^C, ^15^N triple-resonance, z-axis pulsed-field-gradient probes. All NMR data were processed with NMRPipe/NMRDraw (Delaglio et al., 1995). Chemical-shift assignments were obtained from previous work in the lab (Xu et al., 2015). Concentrated RELA-RHD protein was added stepwise to 100 μM ^15^N-TAD protein for the ^15^N HSQC titration. Each sample was concentrated to 300 μL final volume and adjusted to a final concentration of 10% (v/v) D_2_O. ^15^N HSQC titration data were analyzed with CCPNMR (Vranken et al., 2005) with chemical-shift perturbations defined by the equation Δδ_tot_ = [(Δδ^1^H)^2^ + (χ(Δδ^15^N)^2^]^½^ and normalized with the scaling factor χ = 0.15, established from estimates of atom-specific chemical-shift ranges in a protein environment (Farmer et al., 1996).

### ChIP-sequencing analysis

We obtained previously reported ChIP-seq data with antibodies targeting on the RELA/p65 subunit of NF-κB, CLOCK, BMAL1, and their input sample (no specific antibody) of mouse liver treated with LPS from GEO (GSE117488) (Hong et al., 2018). For comparison, we also obtained NF-κB antibody ChIP-seq data and its input sample from mice treated with saline. Each condition contained two biological replicates. The raw fastq reads were aligned to Mus musculus genome assembly (mm10) using Bowtie 2 with sensitive-local option (Langmead and Salzberg, 2012), which can automatically trim the mismatched nucleotide bases (low quality or adaptors) from the end of a read. The biological replicates of the aligned reads were merged together before further analyses. FindPeaks function in the Homer software (Heinz et al., 2010) was deployed to perform peak calling for the NF-κB, CLOCK, and BMAL1 samples with their respective input sample as background control. Significant peaks were declared using the default false discovery rate threshold 0.1%. The comparison of the peaks from NF-κB, CLOCK, and BMAL1 samples were conducted using the mergePeaks function in Homer. Selected peaks were visualized using UCSC genome browser (Karolchik et al., 2003). The peak sequences of ^~^200 bp were used for transcription factor binding motif search using the JASPAR program with a relative profile score threshold of 85% or above (Fornes et al., 2020).

## Abbreviations

NF-κB: nuclear factor kappa B
SCN: suprachiasmatic nucleus
IKK: IκB kinase
IκBα: inhibitor of NF-κB
Luc: luciferase
RHD: Rel homology domain
CRD: C-terminal oscillation regulatory domain
TAD: transactivation domain

## ACKNOWLEDGEMENT

Funding was provided by the National Institute of Neurological Disorders and Stroke (R01 NS054794-06 to ACL and JBH), the National Science Foundation (IOS 1656647 to ACL), the National Cancer Institute (R01 CA149251 to ZW), and American Cancer Society (RSG-13-186-01-CSM to ZW).

---

**Figure S1.**
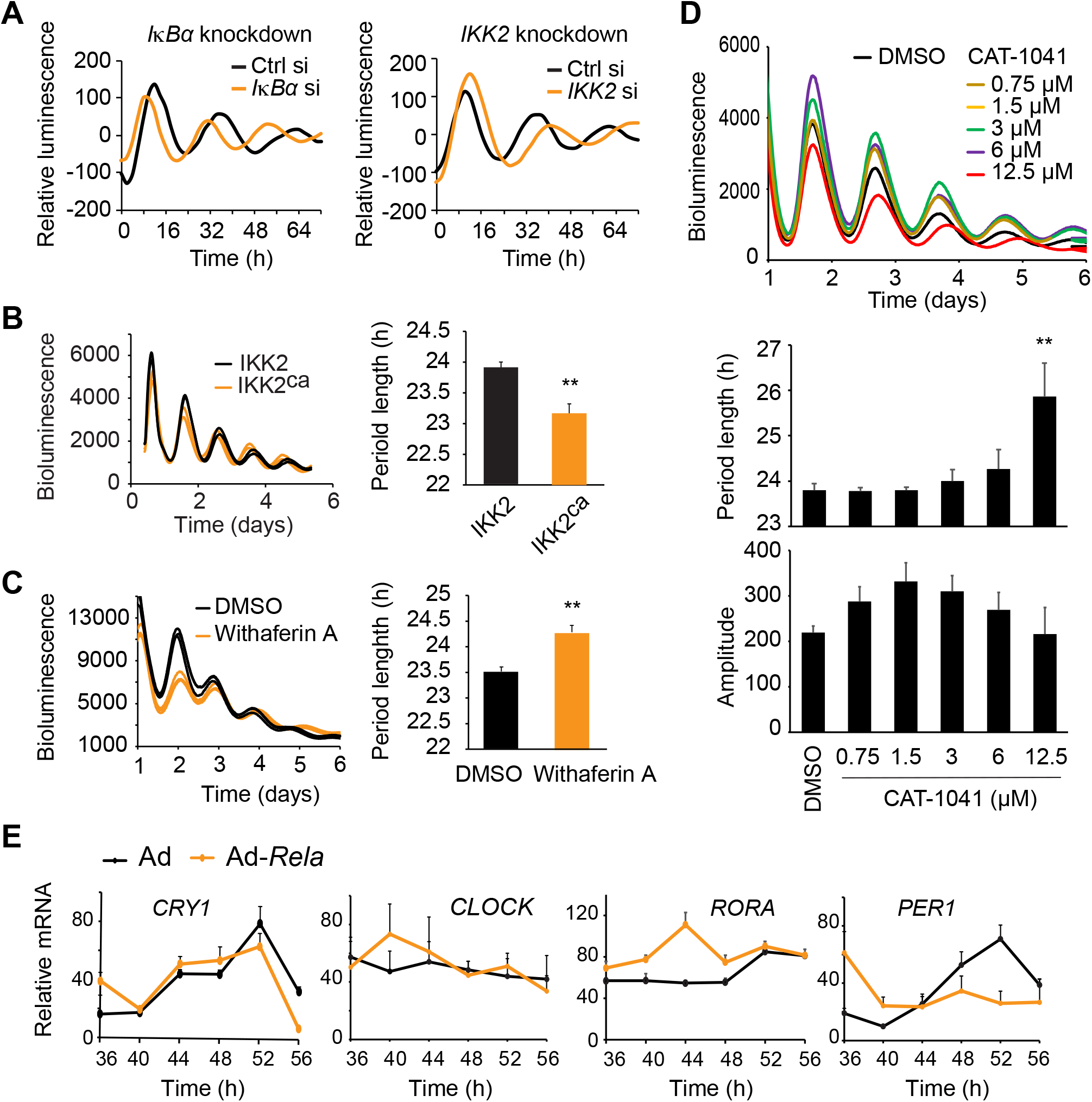
Perturbation of the NF-κB pathway components altered circadian rhythms in U2OS cells harboring the *Per2-dLuc* reporter. **(A)** siRNA-mediated knockdown of *IκBα* and *IKK2* in U2OS cells caused short and long period lengths, respectively. **(B)** Exogenous expression of IKK2-S177/181E (IKK2^ca^) that leads to constitutive acitivation of NF-κB caused short period lengths in U2OS cells. **(C-D)** IKK2 inhibition with chemical inhibitors withaferin A (WA, 1 μM) in (C) or different doses of CAT-1041 in (D) caused long period lengths in U2OS cells. **(E)** NF-κB affected clock gene expression in U2OS cells. Cells were transduced with either adenoviral vector control (Ad) or NF-κB subunit RELA (*Ad-Rela*). The effects of RELA overexpression on the expression patterns of core clock genes were determined by Q-PCR. See Fig. 1 for detail. ** p < 0.01 relative to control.

**Figure S2.**
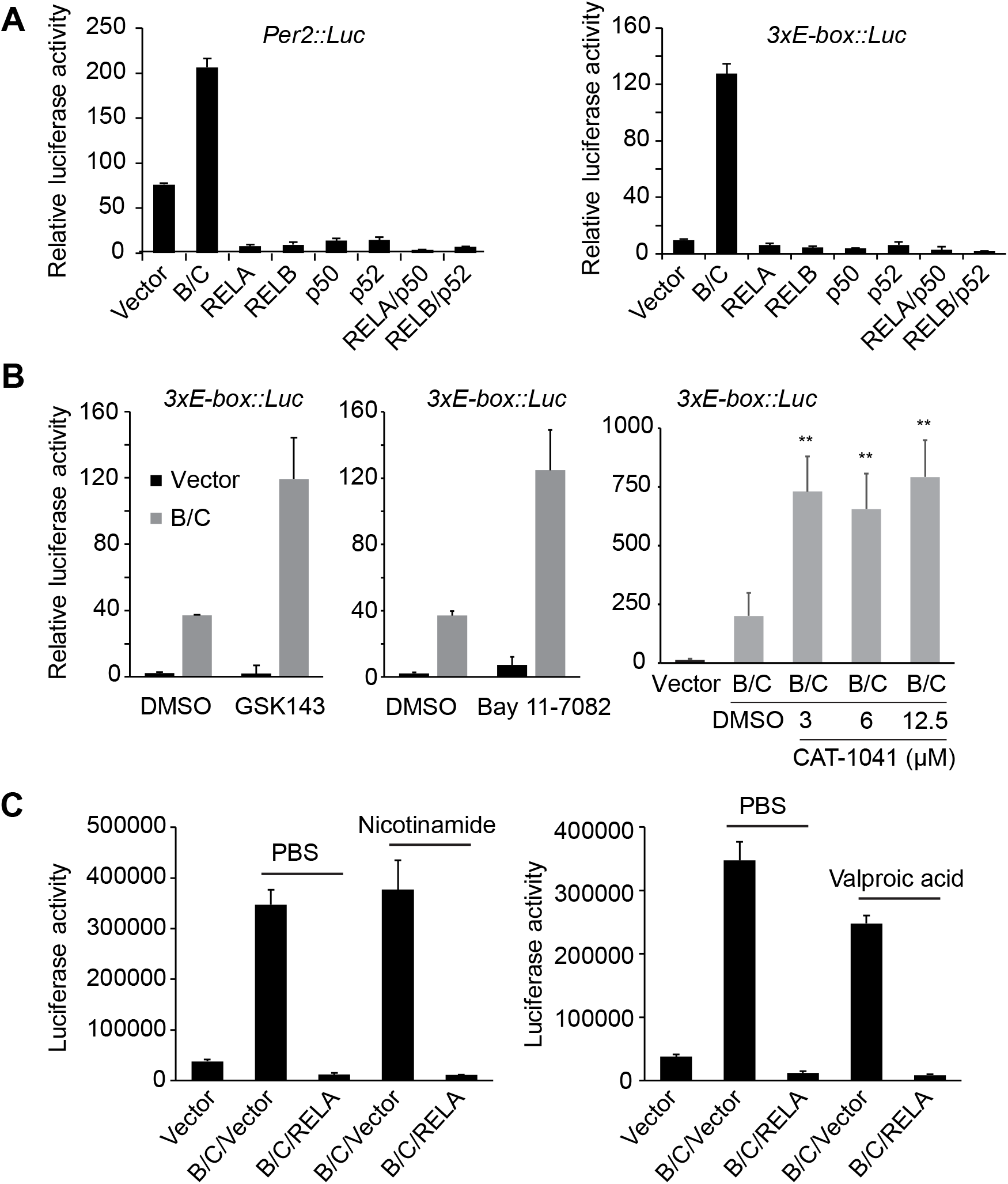
NF-κB affects E-box-mediated transcription. Steady-state luciferase reporter assay in transiently transfected 293T cells. **(A)** NF-κB itself, in the absence of BMAL1/CLOCK (B/C), did not activate the E-box transcription. REAL, RELB, p50 and p52 are NF-κB subunits. **(B)** Inhibition of the endogenous NF-κB by chemical inhibitors (10 μM GSK143, 20 nM Bay 11-7082, or CAT-1041 as indicated) relieved its repression on the E-box-mediated transcription of the luciferase reporter. **(C)** HDAC inhibitors nicotinamide (1 mM) and valproic acid (10 mM) did not affect NF-κB repression of E-box transcription. ** p < 0.01

**Figure S3.**
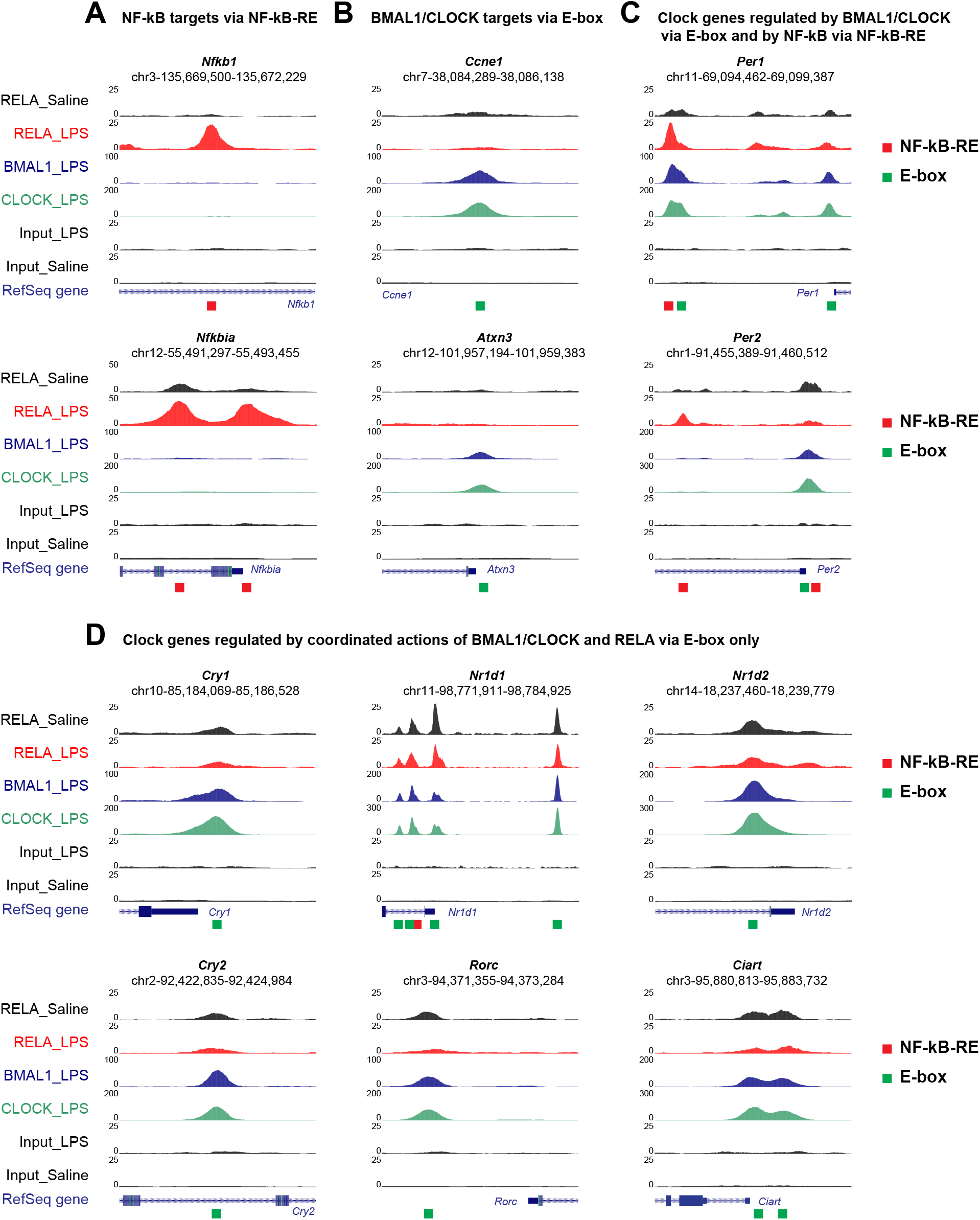
Representative UCSC genome browser images of RELA, BMAL1, and CLOCK ChIP-seq tracks at the genes. ChIP-seq analysis revealed overlapping binding of BMAL1, CLOCK and RELA at the E-boxes of the indicated clock genes. Normalized tag counts are indicated on the Y-axis. The genomic DNA for the region (chromosome # and sequence start and end position) is shown on the top. The predicted motif are shown on the bottom. Green square, E-box. Red square, NF-kB-RE. Motif prediction can be found in Table S1. Note, although motif search for *Nr1d1* predicted an NF-kB-RE site at the second peak, the sequence is located at the peak periphery, not at the peak center (Table S1), and the peak was not LPS inducible, suggesting that NF-kB binding at the second peak is mediated by the two E-boxes at the peak center.

**Figure S4.**
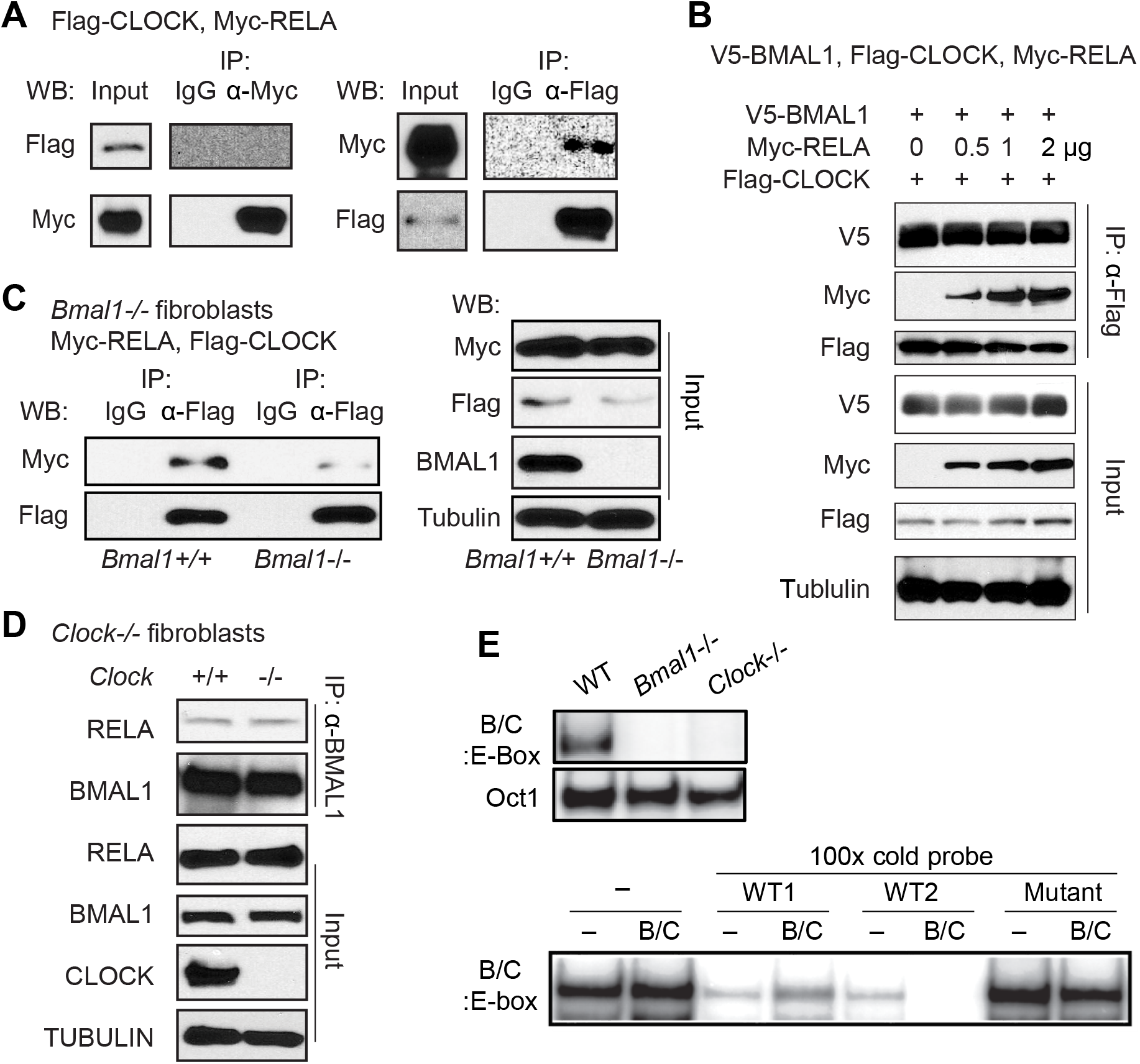
RELA associates with BMAL1 on the E-box. **(A)** Reciprocal co-IP and Western blot to detect the interaction between RELA and CLOCK. Their interaction was not consistently detected in cotransfected 293T cells even after long exposure, indicative of weak interactions. **(B)** Interaction between RELA and CLOCK was readily detected in co-transfected 293T cells in the presence of coexpressed BMAL1. **(C)** The RELA and CLOCK interaction was drastically reduced in *Bmal1*-deficient mouse fibroblasts. **(D)** Co-IP and Western blot detected BMAL1 and RELA interaction in *Clock*-deficient mouse fibroblasts. **(E)** Electrophoretic mobility shift assay (EMSA). The BMAL1/CLOCK (B/C) heterodimer formed a complex with ^32^P-labeled E-box duplex probe (B/C:E-box) in WT cells (left), but not in cells deficient in *Bmal1* or *Clock*, indicating that the BMAL1/CLOCK dimer is required for the BMAL1/- CLOCK:E-box ternary complex formation. The specificity of complex formation was confirmed by completion with a 100-fold excess of unlabeled (cold) wild type probe (WT1, 2), but not with mutated duplex (Mutant).

**Figure S5.**
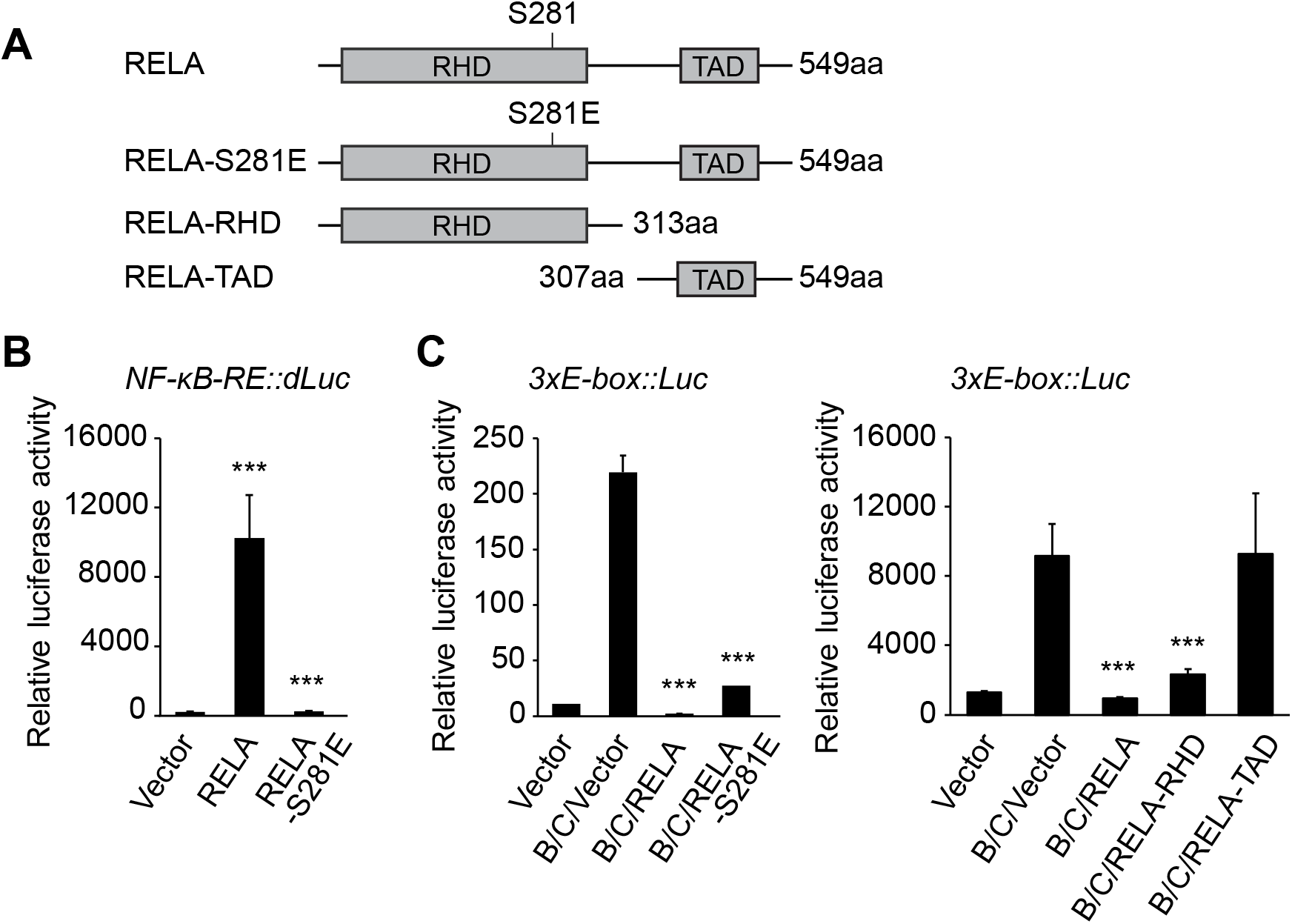
RELA interacts with the BMAL1 C-terminal transactivation domain. **(A)** Schematic diagram of domain structure of RELA. RHD: Rel homology domain. TAD: transactivation domain. Shown are the point mutation and truncation constructs used in the study. **(B-C)** Steady state luciferase reporter assay in transiently transfected 293T cells using the *NF-κB-RE::dLuc* (B) or *3xE-box::Luc* reporter (C). The RELA-S281E mutant failed to activate the *NF-κB-RE::dLuc* reporter, but effectively repressed the *3xE-box::Luc* reporter. The RELA RHD retains the E-box repression activity.

**Table S1.**
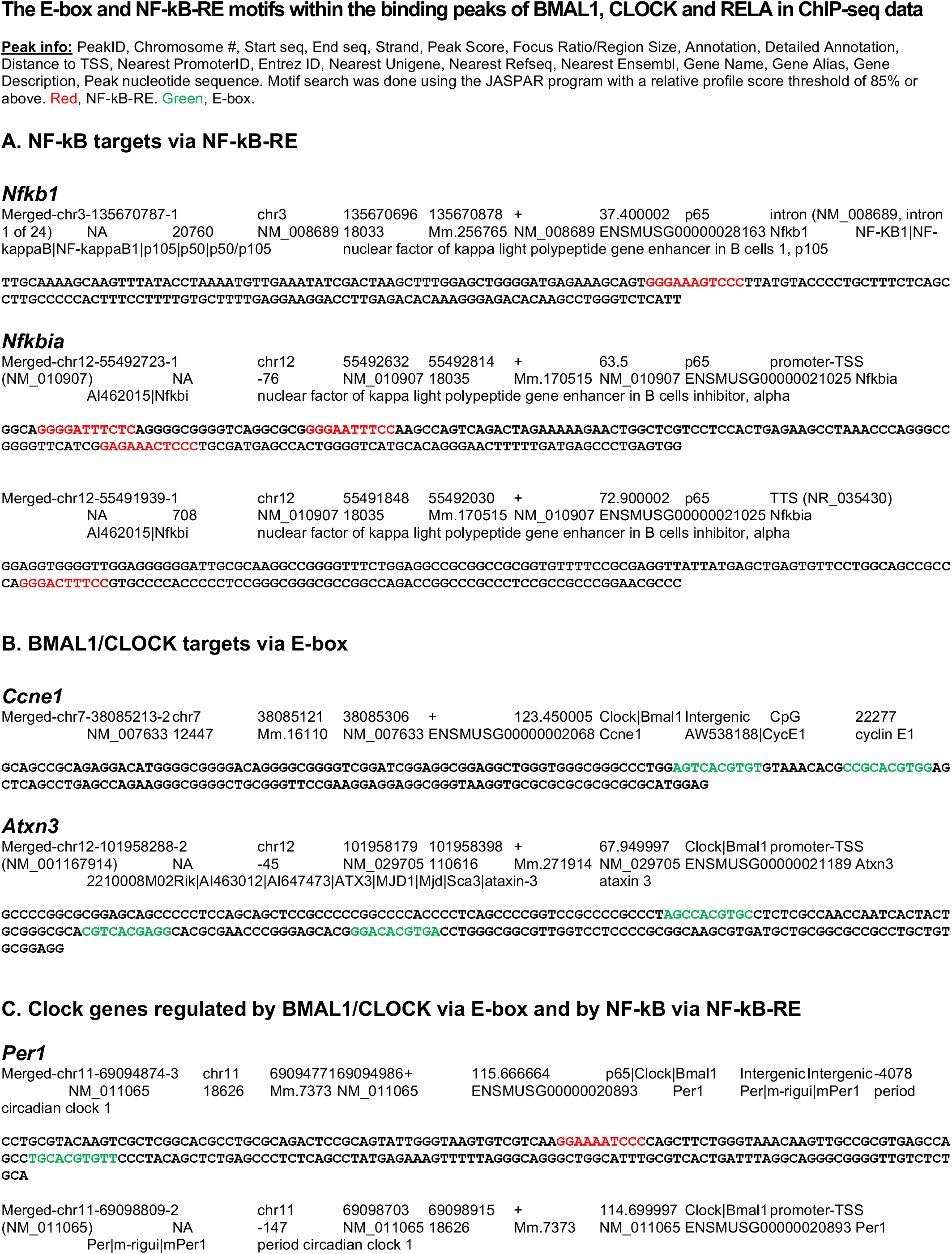
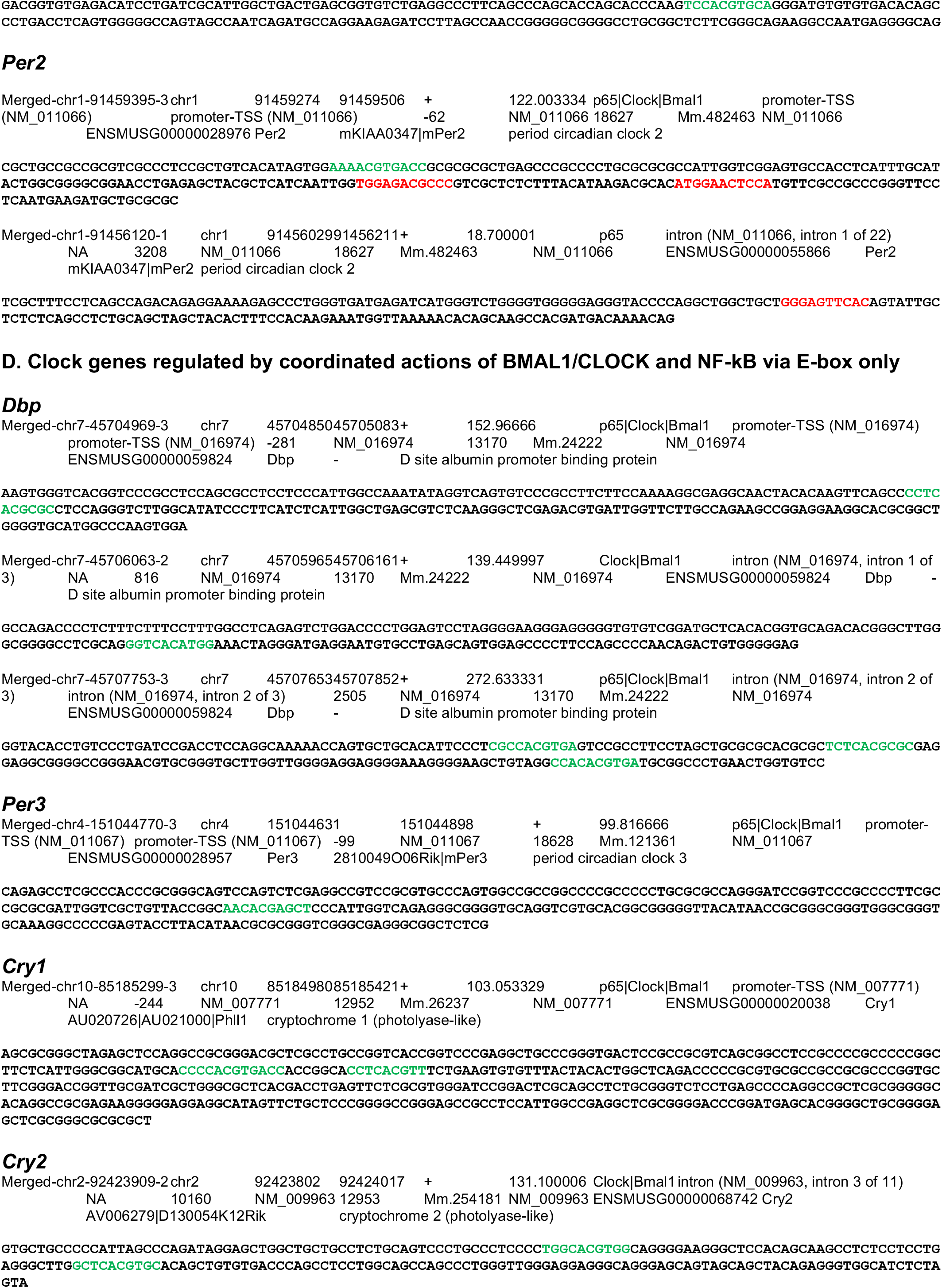
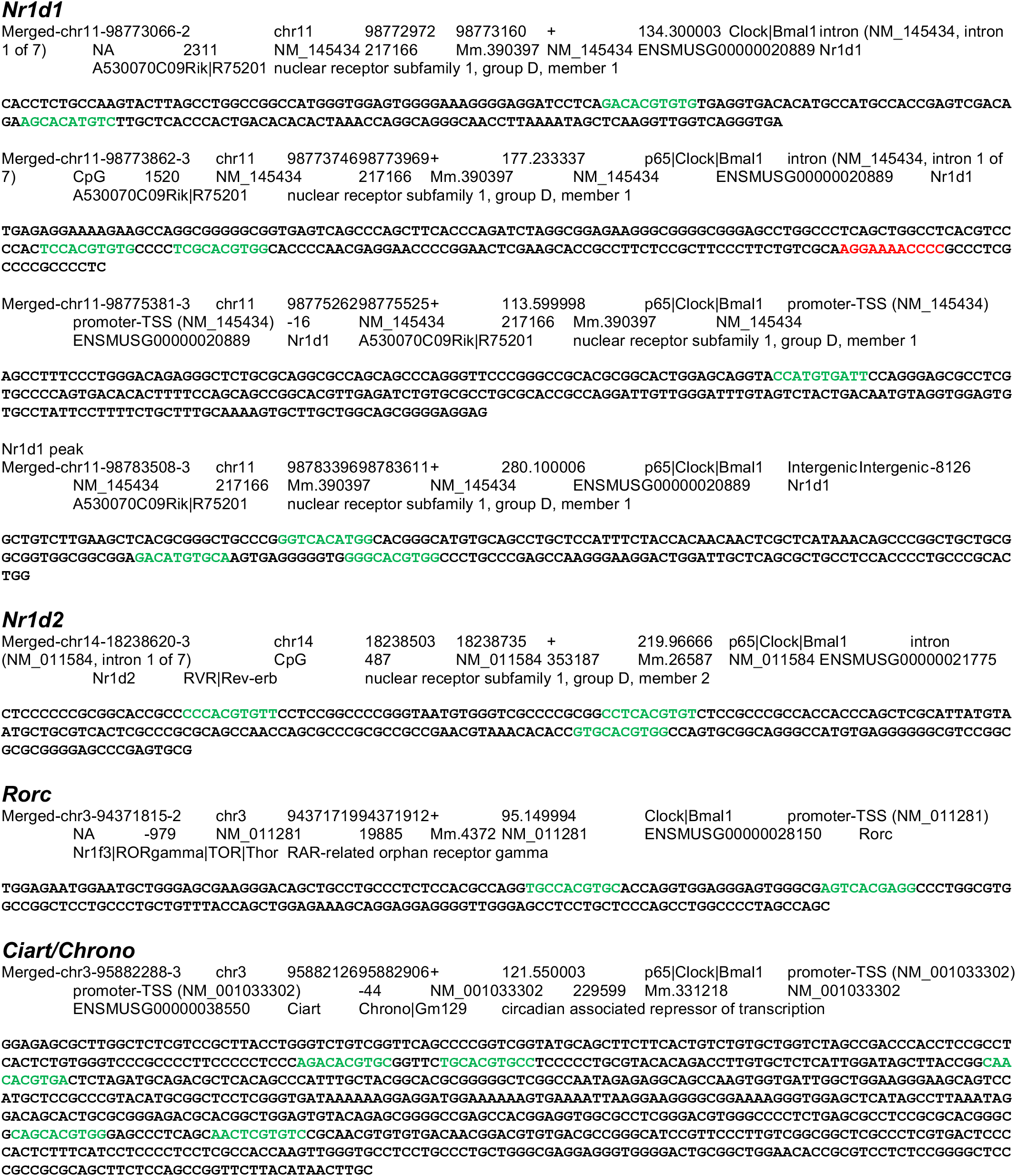

**Table S2.**
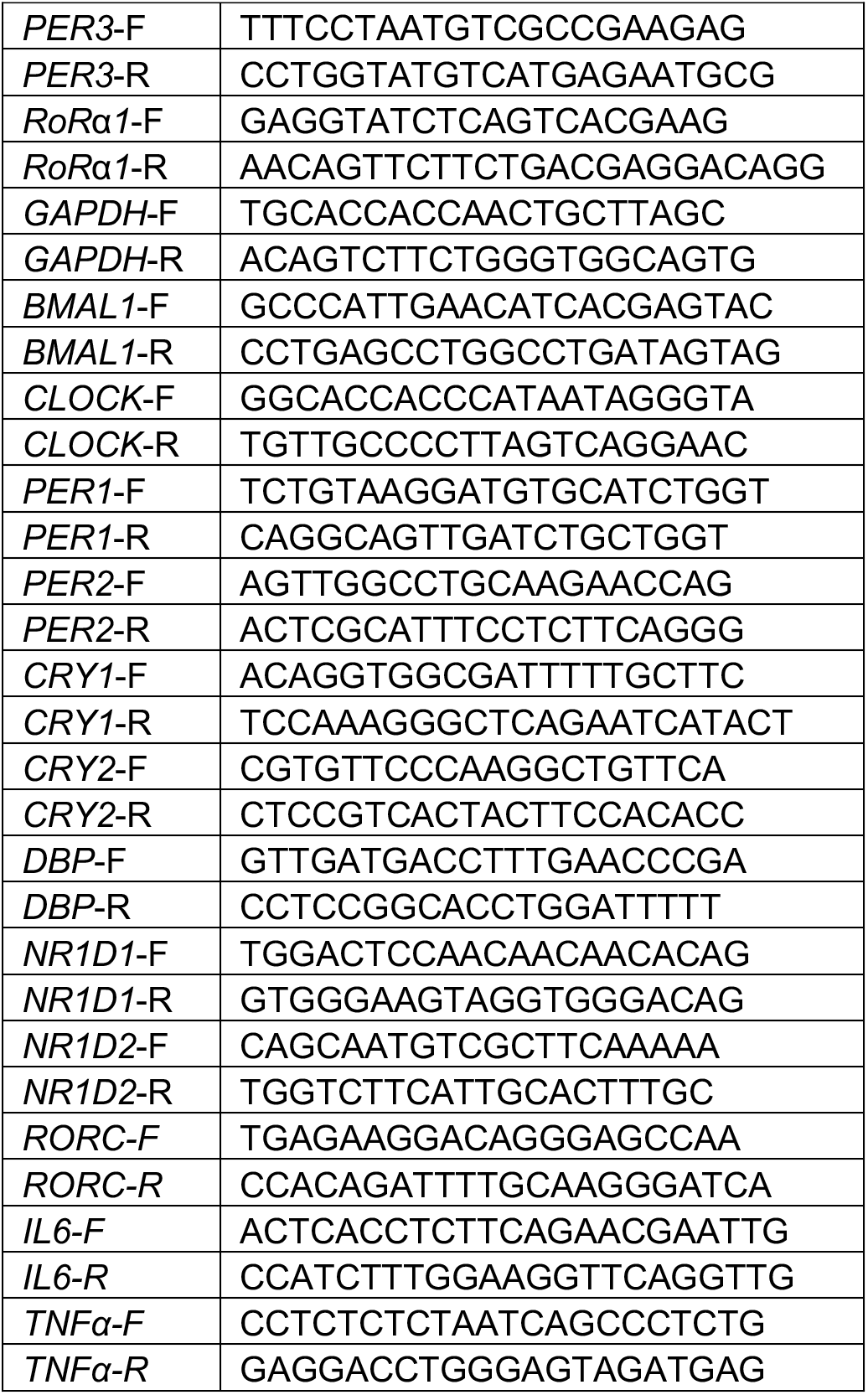
Q-PCR primers for human genes

**Table S3.**
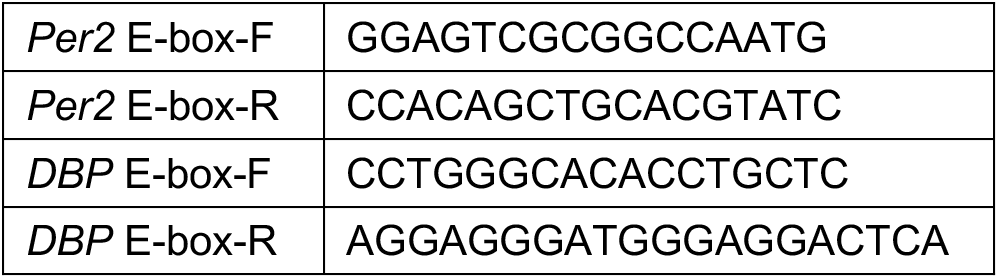
Q-PCR primers for ChIP-PCR

**Table S4.**
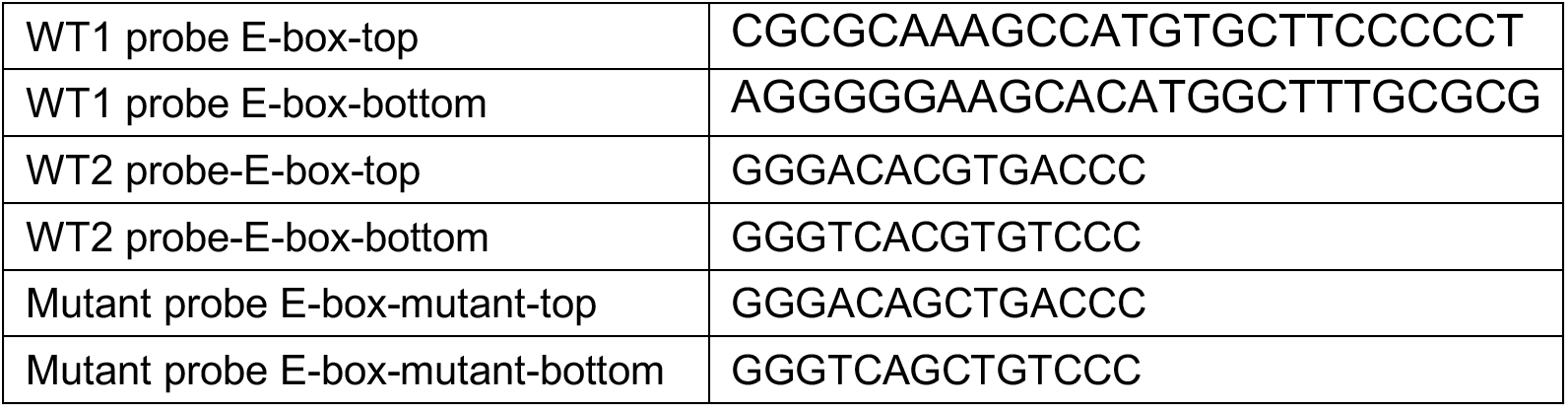
Probe sequences for gel shift

